# Predictive model of transcriptional elongation control identifies trans regulatory factors from chromatin signatures

**DOI:** 10.1101/2022.06.21.496993

**Authors:** Toray S. Akcan, Matthias Heinig

## Abstract

Promoter-proximal Polymerase II (Pol II) pausing is a key rate limiting step for gene expression. DNA and RNA-binding trans-acting factors regulating the extent of pausing have been identified. However, we lack a quantitative model of how interactions of these factors determine pausing, therefore the relative importance of implicated factors is unknown. Moreover, previously unknown regulators might exist. Here we address this gap with a machine learning model that accurately predicts the extent of promoter proximal Pol II pausing from large scale genome and transcriptome binding maps, as well as gene annotation and sequence composition features. We demonstrate high accuracy and generalizability of the model by validation on an independent cell line which reveals the model’s cell line agnostic character. Model interpretation in light of prior knowledge about molecular functions of regulatory factors confirms the interconnection of pausing with other RNA processing steps. Harnessing underlying feature contributions we assess the relative importance of each factor, quantify their predictive effects and systematically identify previously unknown regulators of pausing. We additionally identify 16 previously unknown 7SK ncRNA interacting RNA-binding proteins predictive of pausing. Our work provides a framework to further our understanding of the regulation of the critical early steps in transcriptional elongation.

**Key Points: Please provide 3 bullet points summarizing the manuscript’s contribution to the field:**

- ML model that accurately predicts promoter proximal Pol II pausing from ChIP and eClip-seq data
- Quantification of the interconnection of pausing and other steps of gene regulation
- Identification of novel putative trans regulators of pausing

**GRAPHICAL ABSTRACT:** 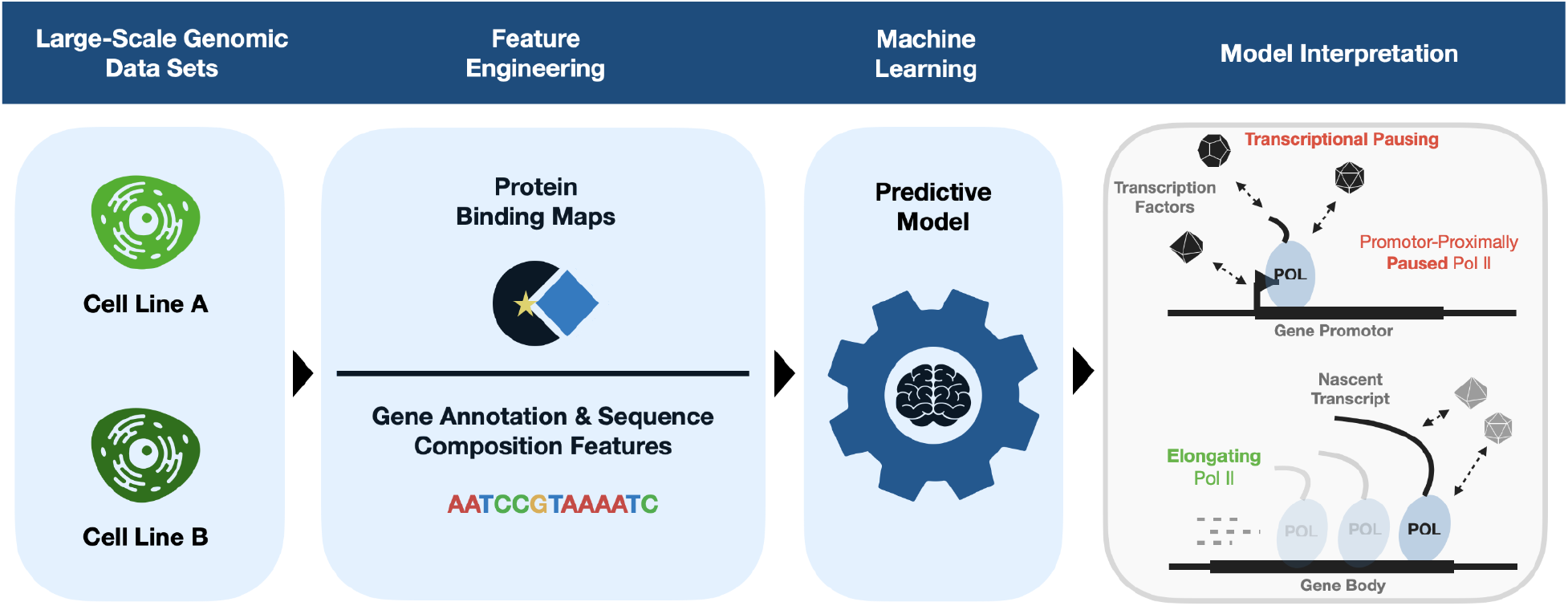

## INTRODUCTION

Transcription of genes is an essential mechanism to maintain cell homeostasis and enable adaptation to changing internal and external stimuli (1, 2). It is tightly regulated by chromatin state and transcription factors (TFs) functioning in a highly coordinated fashion (3). The transcriptional cycle starts with the recruitment of the RNA polymerase into the pre-initiation complex (PIC) (4, 5). During transcription initiation a short fragment of nascent RNA is synthesized. The polymerase is then paused at the promoter before entering into productive elongation upon further regulatory signals or to terminate prematurely (6). This promoter proximal pausing is a key rate limiting step for gene expression as it decides whether a full length transcript will be made or not (7, 8). At equilibrium, paused RNA polymerase accumulates at the promoter since the rate of transcriptional initiation is faster compared to the rates of productive elongation or premature termination (9, 10). In vivo, this accumulation can be observed in assays that monitor nascent transcription such as global run-on sequencing (GRO-seq) (11). Based on this data the equilibrium between transcription initiation and productive elongation, which is decisive for the regulation of gene expression, can be quantified by the pausing index (PI), also known as the travelling ratio (TR) (12–14). It is defined as the ratio of GRO-seq reads in a window around the promoter compared to the rest of the gene body.

Promoter proximal pausing is the default state after transcription initiation (10, 15–17). In addition the duration of pausing is regulated by the interplay of specific factors that either promote pausing or elongation (16). Pause promoting factors include the DSIF complex consisting of Spt5 and Spt4, the negative elongation factor NELF, the 7SK complex, consisting of the most highly expressed non-coding RNA 7SK and proteins such as LARP7 and also specific features of the DNA / RNA sequence (7, 18–22). The most important elongation promoting factor is the positive transcription elongation factor B (PTEFb), which consists of CDK9, CCNK, CCNT1 and CCNT2 (23, 24). Biochemical blocking of P-TEFb showed that its activity is critically important for pause release (25–29). Positive and negative regulators are tightly interlinked. PTEFb is bound by the inactivating 7SK complex and can be released into its active form by BRD4 (30). Once active it phosphorylates regulators of elongation, such as DSIF, as well as other regulators of chromatin state and RNA processing (31). In addition to these direct regulators, pausing is also indirectly regulated by factors that determine transcriptional initiation and transcript processing (32, 33). For example, SRSF2 regulates splicing and has been demonstrated to also determine the duration of pausing (34, 35).

Recruitment of PTEFb to specific promoters through interactions with individual TFs (e.g. NFKB), Mediator, coactivators, and RNA-binding proteins (e.g. DDX2, SRSF2) has been described (34, 36–38). Large scale binding maps of hundreds of RNA binding proteins (RBPs) have recently become available from the ENCODE project (39). Together with the DNA binding maps and GRO-seq these data allow to systematically address several key questions about the regulation of pausing at specific promoters. First, which sequence or protein factors determine the recruitment of regulators to a specific promoter? Second, how do signals from positive and negative regulators translate into the extent of pausing quantitatively?

Here we address these questions by training machine learning models that predict the extent of promoter proximal pausing quantified by the pausing index from large scale genome and transcriptome binding maps as well as gene annotation and sequence composition features. We demonstrate high accuracy and generalizability of the model by validation on an independent cell line and we show that the model can accurately predict differential pausing between cell lines indicating that the model captured general cell type independent rules of pausing regulation. Model interpretation allows for assessing the relative importance of each factor, to quantify their effects and predictive values, and to systematically identify previously unknown regulators of pausing. Grouping of factor contributions by molecular functions confirmed the strong interconnection of pausing and co-transcriptional splicing and other steps of gene expression. We additionally identified 16 previously unknown 7SK interacting RBPs predictive of pausing. These novel pause regulators allow for a systematic and targeted investigation of the regulation of pausing at specific promoters in more detail. Moreover, they provide entry points for experimental manipulation (e.g. with knockdown experiments) to assess their downstream effects on pausing and gene expression in general.

## MATERIALS AND METHODS

### Transcript Annotations (GENCODE)

To engineer gene-centric features of protein binding events and gene annotation and sequence composition features as predictors in our machine learning models we obtained transcript annotations for protein coding genes and non-coding RNAs from the GENCODE (40) database for the hg19 (GrCH37) genome build. We obtained 81745 annotated protein coding transcripts for 20167 genes. Of these transcripts, 30186 (18889 genes) were supported by RefSeq (41) annotations and selected as high confidence transcripts for the analysis. From the annotations we obtained 5-prime, intronic, coding exonic, and 3-prime genomic regions for each transcript which served to capture interpretable binding sites when integrating CHIP-seq and eCLIP-seq data sets (see CHIP-seq data integration & eCLIPseq data integration). HUGO gene nomenclatures (HGNC) (42) from GENCODE were used to further annotate the transcripts with their respective gene symbols.

A set of non-coding transcripts was obtained through appropriate filtering of the GENCODE transcript annotation set for transcripts that were annotated as one of *miscRNA, miRNA, snoRNA, snRNA and lincRNA* which represent miscellaneous, micro, small nucleolar, small nuclear and long intervening RNA biotypes, respectively. These non-coding transcripts were used to engineer features for the machine learning task as well as other downstream analyses especially in the context of the 7SK non-coding RNA (see Identification of 7SK Interacting Poteins). Analogous to the protein coding transcripts the genomic regions (5-prime, intronic, exonic, and 3-prime) of non-coding transcripts were used to create binding site features based on CHIP-seq and eCLIP-seq data sets.

### Transcript Quantifications (RNA-seq)

To ensure that only expressed transcripts are considered we obtained pre-processed transcript quantifications from total RNA-seq experiments from the ENCODE (43, 44) project for the K562 and HepG2 cell lines for the hg19 (GrCH37) genome build. Each experiment had two biological replicates. The obtained transcript expressions were required to have a valid ENSEMBLE (45) ID, to be annotated in the aforementioned GENCODE and RefSeq transcript annotation set, to be expressed (fragments per kilobase million (FPKM) > 0) in both of the replicates. The FPKMs were log10-transformed for downstream analyses. After these filtering steps we considered 16403 (K562) and 16670 (HepG2) of the 30186 protein coding transcripts and 2655 and 1950 non-coding transcripts for the K562 and HepG2 cell line, respectively. The transcript quantifications data sets (tsv-files) were taken from ENCODE experiments ENCSR885DVH (K562) and ENCSR181ZGR (HepG2), with accession numbers of replicated experiments ENCFF424CXV and ENCFF073NHK for the K562 cell line and accession numbers ENCFF205WUQ and ENCFF915JUZ for the HepG2 cell line, respectively.

### Transcription Start Site Annotations (CAGE)

To increase the confidence in the expressed transcripts we further integrated Cap-analysis Gene Expression Data (CAGE) (46) transcription start sites (TSS) for the K562 and HepG2 cell lines. CAGE read counts of the most correlated replicates were aggregated per cell fraction per cell line. Reads were normalized to transcripts per million reads (TPMs). Resulting TSS were then parametrically clustered (47) into CAGE transcription start site clusters (CTSS cluster) with a TPM threshold of 0.1. Singletons with TPM less than 0.1 were excluded. Only transcripts whose transcription start site (TSS) was also the dominant CAGE transcription start site (CTSS) in a cell-type specific CTSS cluster were retained. We thereby were left with 16194 and 16412 protein coding transcripts in the K562 and HepG2 cell line, respectively.

### Quantifying Promoter-Proximal Pol II Pausing (GRO-seq)

We integrated Global-Run-On-sequencing (GRO-seq) (48) data to quantify transcriptional pausing at protein coding genes with the commonly used pausing index (PI) also known as the traveling ratio (12, 26). The PIs served as targets to be predicted in a machine learning task. GRO-seq captures the nascent fragments that build up during the transcriptional cycle and thereby allows to assess Pol II productivity based on the nascent RNA fragment output. As it is commonly done in the field, we have defined the PI as the log2 ratio of GRO-seq read counts (number of 30 bp reads overlapping at each position) at the transcription start site (TSS) to the GRO-seq read signals in the gene body. To optimize the PI definition we have built pausing indices with varying TSS window sizes and chose the window size maximizing the negative correlation of the PI with the corresponding transcript expressions (Pearson’s ρ= −0.68 (K562) and ρ= −0.66 (HepG2); see **Supplementary Figure S1** pausing index optimization). This was motivated by the fact that high PIs, representative of transcriptional pausing, should result in low gene expression profiles and vice versa. This led to a sharp TSS window size of 3bp ranging 1bp up- and downstream of the TSS, while rendering the remaining part of the transcripts as the gene body window. Read lengths of 30bp (K562, GSM1480325) and at least 25bp (HepG2, GSM2428726) ensure that the most frequent Pol II pause site and associated components (49) are covered. Each signal (counts of GRO-seq reads within windows) was then normalized by the respective window size. A pseudocount of 1 read was added to each resulting window for the log2 transformation when building the ratio. The PI was calculated for each of the 16194 and 16412 expressed protein coding transcript in a strand specific manner for the K562 and HepG2 cell line, respectively. Only transcripts which solely contained the DNA base letters (A,T,C,G) along the whole transcript were considered. This further led to the exclusion of 16 and 9 protein coding transcripts in the K562 and HepG2 cell line, respectively. This filtering ensures that we exclude reads that might be erroneously mapped such that we capture the full GRO-seq read signals along the remaining transcripts and thereby obtain comparable signal counts. Overlapping protein coding transcripts were excluded given the fact that corresponding GRO-seq signals can not be uniquely ascribed to a particular transcript and consequently would result in convoluted PI signals. Transcripts which had no GRO-seq signal neither at the TSS nor in the gene body were excluded as well (n=129 in K562; n=196 in HepG2). This has led to the consideration of 8426 and 8260 protein coding transcripts in the K562 and HepG2 cell line, respectively (see **Supplementary Figure S2** for distribution of pausing indices). The corresponding GRO-seq wig-files can be found under GEO accessions GSM1480325 and GSM2428726 for the K562 and HepG2 cell lines, respectively.

### DNA Binding Sites (CHIP-seq)

The integration of chromatin immunoprecipitation sequencing (CHIP-seq) (50) data served to engineer features of gene-centric genomic protein binding events as inputs for the machine learning models. These binding sites for DNA binding proteins (DBPs) were obtained from all available CHIP-seq experiments from the ENCODE project for the K562 and HepG2 cell lines for the hg19 (GrCH37) genome build through corresponding peak-called data sets (bed-files). Perturbation experiments were excluded and only optimal irreproducible discovery rate (IDR) (48, 51) thresholded replicated peaks were considered for downstream analyses to increase the confidence in the obtained binding sites. Experiments with antibodies directly against the factor of interest and newer versioned experiments were prioritized over epitope-tagged and older versioned experiments. We thereby obtained 5041190 (K562) and 4138805 (HepG2) genomic binding sites for 309 (K562) and 211 (HepG2) factors (see **Supplementary Tables S1 & S2** for CHIP-seq factors per cell line) that served for feature engineering purposes (see Feature Engineering). ENCODE CHIP-seq accession numbers for each cell line can be found in **Supplementary Tables S3 & S4**.

### RNA Binding Sites (eCLIP-seq)

Enhanced crosslinking and immunoprecipitation (eCLIP-seq) (52) data served to build gene-centric transcriptomic protein binding features. Binding sites of all RNA-binding proteins (RBPs) from the ENCODE project for the K562 and HepG2 cell lines were obtained for the K562 and HepG2 cell lines for the hg19 (GrCH37) genome build through corresponding peak-called data sets (bed-files). Perturbation experiments were excluded and only optimal IDR thresholded replicated peaks were considered. Newer versioned experiments were prioritized over older versioned experiments. We thereby obtained 409839 (K562) and 435015 (HepG2) transcriptomic binding sites for 120 (K562) and 103 (HepG2) factors (see **Supplementary Table S5 & S6** of eCLIP-seq factors per cell line) for feature engineering (see Feature Engineering). ENCODE eCLIP-seq accession numbers for each cell line can be found in **Supplementary Tables S7 & S8**.

### Identification of 7SK interacting proteins

We filtered the GENCODE transcript annotation data set for all 7SK annotated transcripts to enable the identification of known and novel 7SK binding proteins via observed eCLIP-seq signals on corresponding transcripts and assess their predictive value in the context of transcriptional pausing. In particular, 7SK transcripts which were labeled as pseudo versions were included if they were expressed at least at the median expression level of all expressed non-coding transcripts. Their inclusion was motivated by the idea that factors that also bind these pseudo 7SK transcripts may compete (53) for respective binding sites with factors that bind the non-pseudo version. The set of 7SK binding factors was defined for each cell line as all factors with at least one eCLIP binding site on any of the 7SK transcripts (see **Supplementary Tables S9 & S10**).

### Feature Engineering

For the machine learning task of predicting the gene-wise pausing index of protein-coding genes we engineered features of DNA- and RNA binding events at protein-coding and the closest proximal non-coding transcripts upstream and downstream of the TSS of each protein-coding transcript as well as DNA sequence and annotation features of protein-coding transcripts as predictors for the models. The following features were created:

- transcript length
- strand specification
- chromosome specification
- location on the linear genome
- number of annotated exons
- average exon width
- exon density (ratio of the length of the transcript including introns to the number of exons)
- fraction of exonic sequence (ratio of the length of all exonic sequences to the transcript length)
- GC content of the whole transcript including introns
- Width of CAGE transcription start site cluster (CTSS)
- GC content of CTSS
- distance to most proximal CpG island along with information about the CpG island (length, and features of the sequence: number of CpGs, number of C and G, percentage of CpG, percentage C or G, and ratio of observed to expected CpG)
- binary encoding whether the transcript is a housekeeping gene
- binary encoding of RBP binding events separately for 5’/3’-UTR, introns and coding exons
- binary encoding of DBP binding events separately for 5’/3’-UTR, introns and coding exons excluding Pol II bindings as these are expected to be naturally correlated with the prediction target
- binary encoding of RBP/DBP binding events separately for 5’/3’-UTR, introns and coding exons of the two most TSS proximal non-coding RNAs excluding Polymerase II bindings as these are expected to be naturally correlated with the prediction target

CpG island annotations were taken from the UCSC golden path for the hg19 genome build (cpgIslandExt.txt.gz). Annotations of housekeeping genes were taken from (54). The number of proximal ncRNAs was fixed to two since in combination with CHIP-seq and eCLIP-seq signals on these proximal ncRNAs the feature space would otherwise overgrow the number of genes (and therefore data points in the regression task) which would result in overfitting of the models. Numeric features not in the range [0:1] were rescaled to that range to achieve faster and more accurate model convergences. DNA- and RNA-binding signals went into the model as binary features (binding (1) or non-binding (0)) (see **Supplementary Tables S11 & S12** for number of binding events per factor on individual genomic or transcriptomic regions for each cell line). Distribution of annotation based features for the K562 and HepG2 cell lines can be found in **Supplementary Figures S3 and S4**, respectively. These feature vectors served as a scaffold to build various data matrices for a machine learning regression task based on different feature sub-spaces based on prior domain knowledge as discussed in the next section.

### Feature subsets based on prior knowledge

We stratified the feature space into functionally related sets of proteins in order to characterize the relevance and quantify the importance of pre-, co- or post-transcriptional events in the context of transcriptional pausing. These subsets of binding features of DNA- and RNA-binding factors implicated in specific biological processes were constructed by integrating Gene Ontology (GO) (55, 56) annotations. Functional sets of factors (Chromatin, Initiation, Elongation, Termination, Splicing) were generated based on whether a specific factor was annotated to a biological process (BP) ontology term of any of the following, representative for these major biological functional sets: **Chromatin (**chromosome organization, GO:0051276; chromatin organization, GO:0006325; chromatin remodeling, GO:0006338), **Initiation (**RNA polymerase II preinitiation complex assembly, GO:0051123; transcription initiation from RNA polymerase II promoter, GO:0006367), **Elongation (**transcription elongation from RNA polymerase II promoter, GO:0006368), **Termination (**termination of RNA polymerase II transcription, GO:0006369), **Splicing (**mRNA splicing via spliceosome GO:0045292; regulation of alternative mRNA splicing via spliceosome, GO:0000381) and **Processing (**mRNA export from nucleus, GO:0006406; mRNA 3’-end processing, GO:0031124). The set of Elongation factors was further extended by pause regulatory factors from the literature (16, 57, 58) if not already included in the GO-derived factor set **Elongation**. These were super elongation complex (SEC) factors CCNT1, CCNT2, ELL, ELL2, ELL3, AFF1, AFF4, MLLT1, MLLT3, established pausing factors NELFA, NELFB, NELFCD, NELFE, SUPT4H1, SUPT5H, SUPT6H, SUPT16H, BRD4, MYC, TAF1, TBP, PAF1, and CDK9 (P-TEFB), as well as 7SK ncRNA pause mediator complex binding factors LARP7, HEXIM1, HEXIM2 and MEPCE (see also **Supplementary Table S13**). However, we could only consider a subset (n=19) of all established pausing factors, which were assayed in the CHIP-seq and eCLIP-seq experiments. The final **Elongation** factor set thus contained POLR2A, POLR2B, POLR2G, POLR2H, MLLT1, SUPT5H, GTF2F1, BRD4, WDR43, NCBP2, HNRNPU, LARP7, MYC, TAF1, TBP, AFF1, EZH2, PAF1 and SSRP1. However, Polymerase associated factors (POLR2A, POLR2B, POLR2G, POLR2H) were excluded since these are expected to correlate with the pausing signal. A set of 7SK binding proteins derived from binding sites observed in the eCLIP-seq data was generated to quantitatively assess the importance of unknown or less well established 7SK associated factors (see 7SK non-coding RNA or **Supplementary Table S4** of 7SK binding factors per cell line). A set representative of general pausing associated factors was generated by forming the union of the Elongation and 7SK associated factor set (**Elongation+7SK**). For a list of factors in each resulting functional factor set per cell line see **Supplementary Tables S14 & S15**.

Each resulting factor set was further stratified into sequence-specific and non-sequence-specific binders. The Molecular Signatures Database (MSigDB) (59, 60), a collection of annotated gene sets, the Catalog of Inferred Sequence Binding Preferences (CIS-BP) (61), a library of transcription factors and their binding motifs and the Homo sapiens comprehensive model collection (HOCOMOCO) (62), a collection of transcription factor binding models for human and mouse via large-scale ChIP-seq analysis based on binding motifs, were queried to identify sequence specific factors (see **Supplementary Tables S16 & S17**).

The feature vector space of binding events was then accordingly grouped by these factor sets (see **Supplementary Table S18** of factor presence in feature subspaces) to form different feature matrices, always accompanied with DNA sequence and annotation features of protein coding genes. These feature matrices based on prior domain knowledge, 7SK ncRNA associations and sequence-specificity served to build an array of predictive models based on features with a defined biological function. For a baseline comparison of model performances we have further built 100 random models which randomize over the number of factors, the factors itself and their binding patterns. The binding patterns were randomized according to the observed binding proportions.

### Model Training

Models of transcriptional pausing were obtained by training Extreme Gradient Boosting Tree (XGB) regressors to predict the pausing index with each of the feature subsets (see Feature Stratification). Apart from 5-fold cross validating the models on the training cell line, we applied trained models to the completely independent test data sets from the other cell line. This provided us with an unbiased estimate of the model performances as trained models have neither seen the genes target distribution nor the specific feature distributions of the other cell type. To enable such validation procedure each feature matrix was reduced to features common to both cell lines. We refer to these as the *synchronised* models as compared to the *individual* models which on the other hand incorporated all available features specific to a cell type. In the case of individual models 50% of the available data points were held out prior to training as an independent test data set. Though this set is not from an independent cell line as is the case with the synchronised models, it still provides an unbiased model performance estimate as trained models have also not seen any of the data points from the hold out test data set.

Regression with squared loss was chosen for the learning objective. The coefficient of determination (R-squared, R^2^) was used as the evaluation metric to compare and evaluate trained models. See **Supplementary Table S19** for hyperparameter specification and the zenodo repository for R-Data structures with all model matrices (model.matrices.RDS).

### Feature scoring

Shapley additive explanations (SHAP) (63, 64) were used as a scoring metric for feature contributions. SHAP is a game theoretic approach to explain the output of any machine learning model. In contrast to the well known variable importance metric it is able to show the positive or negative relationship for each feature with the target. As opposed to most feature importance metrics that average over all genes, each gene receives its own set of SHAP values, greatly enhancing the prediction transparency. SHAP values are additive and allow to aggregate over contributions of subsets of features which enabled us to capture contributions of binding features per protein and subsequently group these proteins into sets of positive and negative regulatory factors. For instance, we obtain contribution scores for a transcription factor binding on the 5’UTR, exons, introns and 3’UTR on the genome and transcriptome as identified by CHIP-seq and eCLIP-seq, respectively. We derived total factor contributions by aggregating the SHAP scores per factor over each gene region which enabled us to identify specific pause regulatory factors by selecting factors with high effect sizes.

## RESULTS

### Predictive Models of Transcriptional Pausing

The transitioning of promoter-proximally paused Pol II (Fig. 1A, promoter-proximally paused Pol II) into its elongating phase of nascent RNA synthesis (Fig. 1A, elongating Pol II) is regulated by trans-acting protein co-factors as well as cis-regulatory DNA and RNA sequence features (16, 18) which we refer to as chromatin signatures.

**Figure 1:**
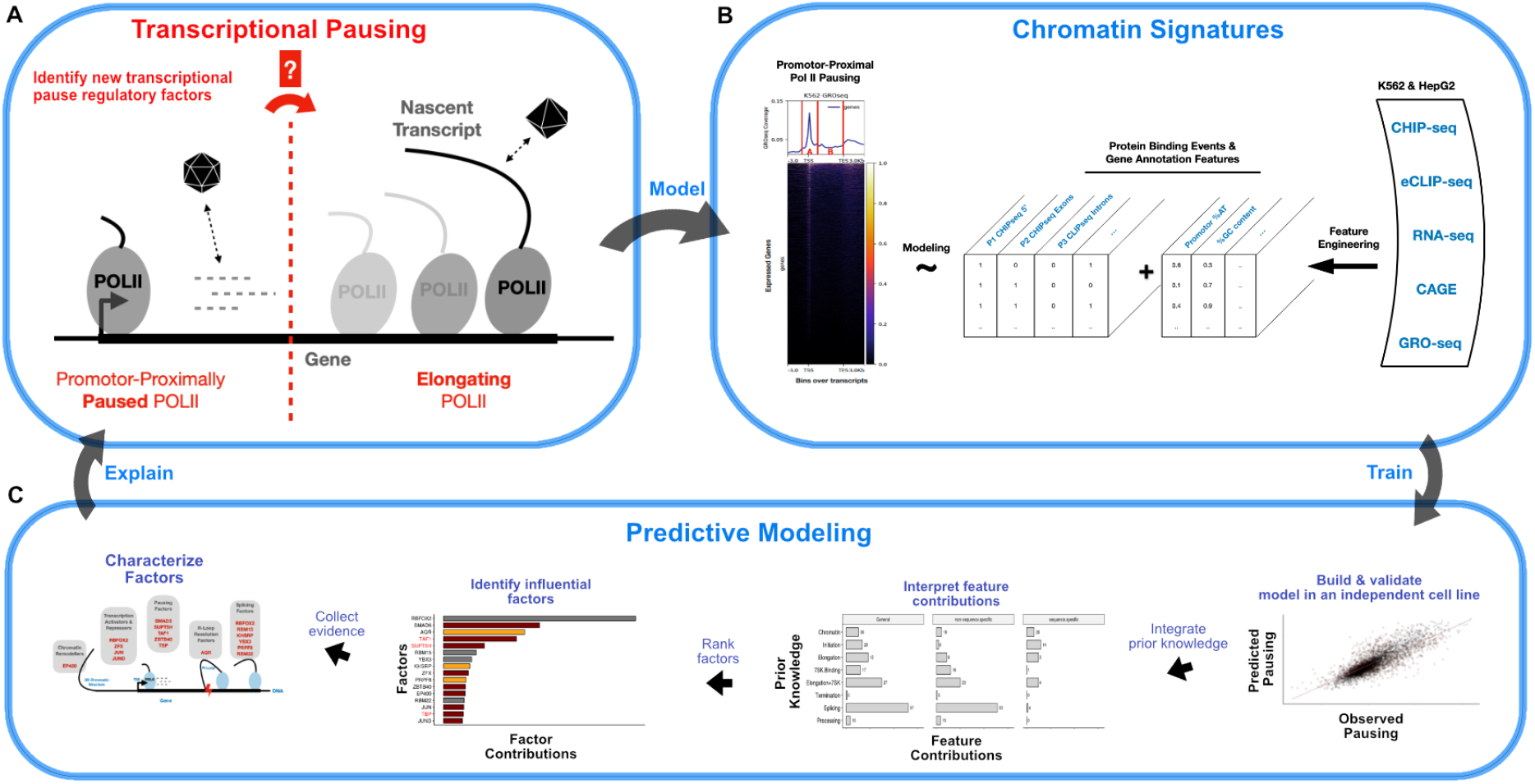
**(A)** Central question as to which specific factors are implicated in the transitioning of promoter-proximally paused Polymerase II into its elongating phase of nascent RNA synthesis. **(B)** Integration of large-scale genomic data sets to build the chromatin context of transcriptional pausing (A) with protein binding events and gene annotation and sequence composition features for the prediction task of promoter-proximal pausing of the Polymerase II quantified by relating GRO-seq read densities at the TSS to GRO-seq read densities in the gene body **(C)** Machine learning approach to predict promoter-proximal Pol II pausing with chromatin signatures (B), followed by the integration of prior knowledge and selection of factors as modulators of promoter-proximal Pol II pausing.

For the identification of such specific regulatory chromatin signatures we used large-scale genomic and transcriptomic protein binding maps from ENCODE and compiled gene annotation and sequence composition features. We then followed a systematic machine learning approach to predict the degree of transcriptional pausing at protein coding genes (Fig. 1B) through the integration of these chromatin signatures in a regression model with Extreme Gradient Boosting trees (XGB) with the potential to reveal explanatory factors (Fig. 1C).

To facilitate the validation in independent cell lines we obtained relevant data sets for two different cell lines (K562 and HepG2). The prediction target was defined as the gene-wise *pausing index* (see Materials & Methods; see **Supplementary Figure S2** for pausing index distributions). It quantifies the degree to which a gene is paused (high pausing index) or elongated (low pausing index). To construct the feature matrix of predictors as input for our models we systematically integrated genome-wide CHIP-seq (see Materials & Methods) and eCLIP-seq (see Materials & Methods) data from the ENCODE project, providing DNA and RNA binding sites on the genome and transcriptome respectively (see **Supplementary Tables S11 & S12**). Gene-centric annotation and composition features were mainly engineered based on GENCODE transcript annotations (see Materials & Methods, **Supplementary Figures S3 and S4**). CAGE transcription start sites were integrated (see Materials & Methods) to define high confidence TSS and further validate the expression of transcripts. We thereby obtained a total of 2503 features of 2485 DNA & RNA binding and 18 gene annotation features in the K562 cell line and 1832 features of 1814 DNA & RNA binding and 18 gene annotation features in the HepG2 cell line. We then trained an Extreme Gradient Boosting Tree regressor (see Materials & Methods and **Supplementary Table S20**) to predict the pausing index of protein coding genes (n=8426 in K562) with high accuracy and explain up to 68% of the observed variance (R^2^~0.68 on 50% hold-out test data set, K562) of the pausing index (Fig. 2A).

**Figure 2:**
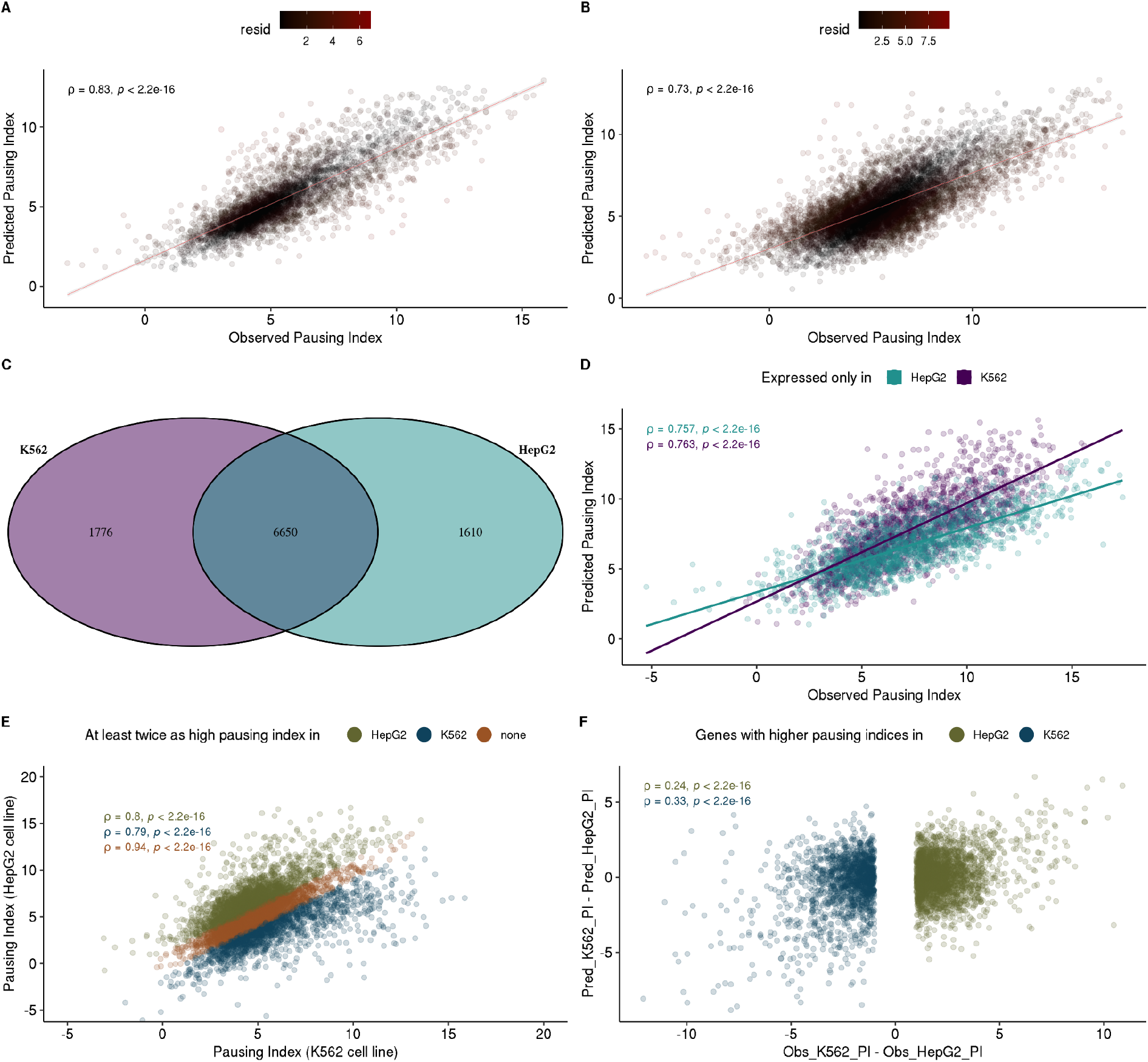
(**A**)Observed vs. predicted pausing indices (log2 scale) of a 5-fold cross-validated and regularized XGB regression model in the K562 cell line applied to an independent 50% hold-out test dataset from the same cell line taken prior to training. Pearson’s correlation coefficient rho (ρ) with the associated p-value is depicted in the upper left. The residual regression error is colored in red (see legend *resid*). **(B**)Observed vs. predicted pausing indices of a 5-fold cross-validated and regularized XGB regression model in the K562 cell line applied to the independent test dataset from the cross cell line (HepG2). Model was trained with features common to both cell lines. Pearson’s correlation coefficient rho (ρ) with the associated p-value is depicted in the upper left. The residual regression error is colored in red (see legend *resid*). **(C)** Venn diagram of transcripts between cell lines. **(D)** Observed vs. predicted pausing indices of a 5-fold cross-validated and regularized XGB regression model from each cell line applied to data of genes exclusively expressed in the cross cell line. Pearson’s correlation coefficient rho (ρ) with the associated p-values are depicted in the upper left. **(E)** Observed pausing indices from the K562 vs. HepG2 cell line. Transcripts with at least a 2-fold higher pausing index in one but not the other cell line are colored either green (HepG2 specific transcripts) or blue (K562 specific transcripts). Transcripts with similar pausing indices (less than a 2-fold change) in both cell lines, thus not specific to any of the cell lines, are colored in orange. Pearson’s correlation coefficients (ρ) for each of the groups with associated p-values are depicted in the upper left. **(F)** Observed pausing index differences between cell lines against differences of predicted pausing indices obtained from models trained in each cell line and applied to data from the cross cell line. Models were trained on features common to both cell lines. Differences are shown for genes which showed a 2-fold change between cell lines as identified in E).

The model performances can be further evaluated through **1)** the application of a model trained on one cell line and applied to the full data of the other cell line (Fig. 2B) **2)** the application of a model trained on one cell line and applied to genes that are only expressed in the other cell line (Fig. 2D) and **3)** the application of a model trained on one cell line and applied to genes present in both cell lines with significantly different pausing indices representing extreme observation specific to the other cell line (Fig. 2F). See **Supplementary Figure S5** for figure 2 analog of model performances of a model trained on the HepG2 cell line and validated on the K562 cell line.

The predictive power and generalizability of the model was supported by the high prediction performance on the independent cross-cell type test data set (Fig. 2B, performance on HepG2 data of K562 model) in which it was still able to explain up to 53% of the variance. The decreased model performance with an R^2^ of 0.53 as compared to 0.68 (Fig. 2A) is likely due to the reduced amount of features that are available in the HepG2 cell line (39% of all features (n=987) of n=2503 features available in the K562 cell line).

A good performance in the cross cell type prediction task (Fig. 2B) can have two reasons: **1)** the model captures the signal of ubiquitously expressed genes which are similar between cell types, as might be the case with housekeeping genes, or **2)** it learned general rules that would also allow for predicting cell type specific pausing indices from cell type specific chromatin signatures. To distinguish these scenarios we identified the sets of exclusively expressed genes (Fig. 2C) and assessed the performances of models trained on one of the cell lines on the genes exclusively expressed in the other cell line (Fig. 2D). The K562 model was able to explain up to 57% and the HepG2 model up to 58% of the observed variance in the pausing indices in the HepG2 and K562 cell line respectively.

We further validated that our model can also identify quantitative changes on transcripts which showed differential (fold change >= 2) cell type specific distributions of the pausing indices. For these sets of transcripts (Fig. 2E, blue, green) we evaluated the concordance of observed pausing index differences between the cell lines against the differences of predictions of the pausing indices using models trained in one of the cell lines and applied to data in the other cell line (Fig. 2F). Although we can recognize a substantial decrease in model performances with a correlation of 0.24 (Fig. 2F, HepG2 specific pausing indices; green) as compared to 0.73 for the prediction on the entire HepG2 cell type data (Fig 2B) or 0.76 on HepG2 cell type specific genes (Fig 2D), the model still maintains predictive power for extreme observations of pausing indices specific to the cross cell type which further underlines the ability of the model to generalize to other cell lines.

Given the high predictive power of the obtained model not only on intra-cell type holdout test data sets (Fig. 2A), the inter-cell type test data set (Fig. 2B) as well as its ability to predict pausing indices of cell type specific genes (Fig. 2D) and cell type specific differential pausing indices (Fig. 2F), we concluded that our model captured general rules of pausing regulation independent of the cell type and that the underlying chromatin signatures of the models would have sufficient discriminatory power to explain the observed variance in the pausing index. We thus continued with downstream feature interpretation and selection approaches to suggest potentially novel regulators of transcriptional pausing. Downstream analyses were performed on data from the K562 cell line due to the increased amount of features available.

### Contribution of individual transcript processing steps to the prediction of pausing

We next aimed to gain a mechanistic understanding of the underlying predictive contributions. To measure the contributions of model features we have used Shapley Additive Explanations (SHAP) (64, 65) as a feature scoring metric (see Materials & Methods) which captures the directional contribution of each model feature specifically for each gene on the target variable. A model feature may increase or decrease the pausing index or exert no effect at all depending on the factors relevance for pausing and their interaction with other features of each gene (Fig. 3A). Their combined effects converge in predicted pausing indices which in turn represent the average output whether a gene is paused or not.

**Figure 3.**
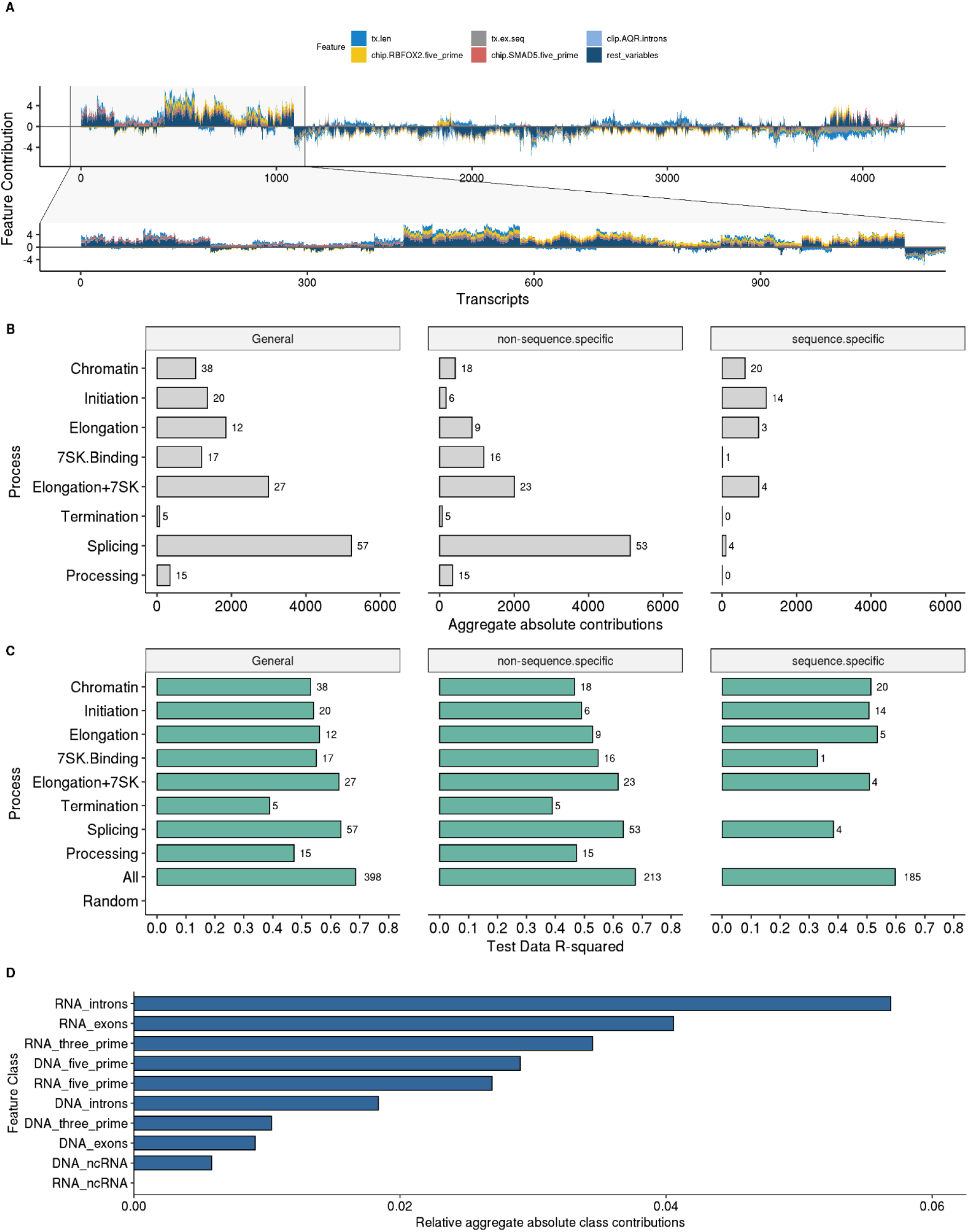
**(A)** Individual feature contributions (y-axis; only top 5 individual features colored and remaining aggregated in *rest_variables*) on each transcript (x-axis) with a sample zoom-in on a subset of transcripts for better visual investigation **(B)** Aggregate absolute contributions of factor classes based on prior knowledge, further divided by sequence and non-sequence specific binding factors. The process *Processing* refers to mRNA polyadenylation and export from the nucleus. **(C)** R^2^ performances of individual models of factor classes based on prior knowledge on 50% holdout test data set **(D)** Aggregate absolute contributions of factors based on their binding modes.

Because transcriptional pausing is connected with other steps of gene expression from chromatin organization (66–68), transcription initiation (8, 49, 69), to splicing (32, 70, 71) and post-transcriptional transcript processing (33, 72, 73), we assessed the regulators of these pre-, co- or post-transcriptional events according to their importances in predicting pausing. To that end we have generated sets of regulators (see Methods and **Supplementary Tables S14 & S15**) representative of specific RNA processing events (*Chromatin, Initiation, Elongation, Splicing, Termination, Processing*) based on Gene Ontology (GO) annotations. The *Elongation* factor set was further extended by established pausing factors from the literature. The 7SK non-coding RNA complex is a key regulator auf pausing (34, 74–78). To assess the role of RNA binding proteins participating in the 7SK complex for pausing, we additional built a set of factors that bind the 7SK ncRNA in the eCLIP-seq datasets (see Methods and **Supplementary Tables S9 & S10** for 7SK binding factors per cell line). This set included the well known 7SK binder LARP7, the pausing related regulator AQR previously not associated with the 7SK as well as the following factors not previously associated with pausing: SSB (LARP3), HNRNPK, DGCR8, PCBP1, ATF, ZNF800, XRCC6, NCBP2, SBDS, YWHAG, GRWD1, ZNF622, SRSF7, TARDBP and BUD13. A set consisting of the union of Elongation and 7SK associated factors was generated as well (*Elongation+7SK*). All sets of regulators were further stratified into known sequence-specific and non-sequence-specific binders (see **Supplementary Tables S16 & S17**) in order to assess the relevance of sequence specific binding events. For each factor in the resulting functional set of regulators we aggregated their feature contributions (Fig. 3A) per functional process (Fig. 3B).

Splicing factors had the highest contributions followed by elongation and 7SK binding proteins. This strongly supported the intricate connection to co-transcriptional splicing events (35, 70, 79) and strengthened the role of the newly identified 7SK binding proteins as transcriptional pause regulatory factors. The *Elongation* factor set of established pausing factors served as a validation of our approach.

We next asked how models would perform if they are trained exclusively on the features defined by each of the previously defined sets of regulators. For a baseline comparison models were also trained on randomized input data (see Materials & Methods). Figure 3C shows the model performances (R^2^ values) for each of the feature subspaces of cross-validated models in the K562 cell line on the independent 50% holdout test data sets (see also **Supplementary Table S20** for all model results). In general all models perform reasonably well relative to the number of features they incorporate. As an example the splicing factor based model (*Splicing*) incorporates only 14% (n=57) of all available factors yet performs almost equally well as the full model (*All*) incorporating all available factors (n=398). Likewise, the *Initiation* model considers only about half the number of factors than the chromatin associated model (*Chromatin*) yet performs slightly better (R^2^ of 0.54 vs. 0.53).

As expected, the 7SK ncRNA associated factor model (*7SK*.*Binding*) and the model with previously established pausing factors (*Elongation*) perform very well despite the low number of factors considered in those models. The predictive power of pausing/elongation factors becomes further evident when we consider the model of the union of 7SK and established elongation factors (*Elongation+7SK*) which outperforms (R^2^ 0.62) each individual factor set alone (*7SK*.*Binding*: R^2^ 0.55, *Elongation*: R^2^ 0.56) and performs almost equally well as the full model (R^2^ 0.62 vs. 0.68). This result highlights the relevance of the novel set of 7SK binders identified by protein-RNA interactions as putative pause regulators. Taken together, the majority of factor sets show high predictive power relative to the number of factors they incorporate but their performances should not be compared to each other due to the variable amount of factors considered in the models. Their predictive value demonstrates the interconnectedness of underlying processes with the transcriptional pausing outcome. It further supported and strengthened the role of the 7SK ncRNA as a transcriptional pause mediator complex and allowed us to suggest the factors from the set of 7SK associated factors (*7SK*.*Binding*) (see **Supplementary Tables S9 & S10**) as additional 7SK ncRNA binding proteins to be implicated in the regulation of pausing based on their predictive value.

We next asked whether protein-DNA or protein-RNA binding events contributed to the explanatory power of the models. We found that the individual contributions of RNA binding events are generally higher than those of DNA binding events (Fig. 3D). Investigating the contributions of factors by their functional classes within the highest ranked class (RNA introns) (see **Supplementary Figure S7 & S8**) reveals that splicing factors are enriched for RNA intron binding sites (Fisher’s exact test, one-sided (greater), p = 0.034, odds ratio 4.45, confidence interval [1.11;Inf] in K562 and p=0.032, odds ratio 7.1 [1.15;Inf] in HepG2). The high contributions of genomic binding events on the 5’ region of transcripts (Fig 3D, DNA_five_prime) are in line with observed genomic five prime modulated transcriptional pause states (80).

Overall the results for the HepG2 cell line are very similar and support the conclusions (**Supplementary Figure S6)**. Although gene annotation and composition features account for 26% of all feature contributions (see **Supplementary Figure S9-12**) they are static in their nature and cannot explain variation of pausing between cell lines. Therefore, we focus the discussion on individual proteins and their binding events as they are dynamic between cell lines.

### Modulators of Transcriptional Pausing

Based on our model, we aimed to identify specific pause regulatory factors. To obtain a ranking of the importance of individual DNA- and RNA-binding factors for predicting Pol II pausing, we aggregated the SHAP contributions (see **Supplementary Figure S13 & S14** for individual feature contributions per cell line) into a single contribution score per factor and selected the minimal set of most influential factors (16 out of 398) that makes up 50% of all feature contributions (Fig. 4A). Established pausing factors from the literature (Fig. 4A, highlighted in red) are ranked among these top influential factors, validating our factor ranking approach. Three factors not primarily related to pausing were ranked higher than the established pausing factors and are potential novel modulators of pausing with at least the effect size of the established factors. However, all other factors have similarly high contributions and can be considered almost equally important.

**Figure 4.**
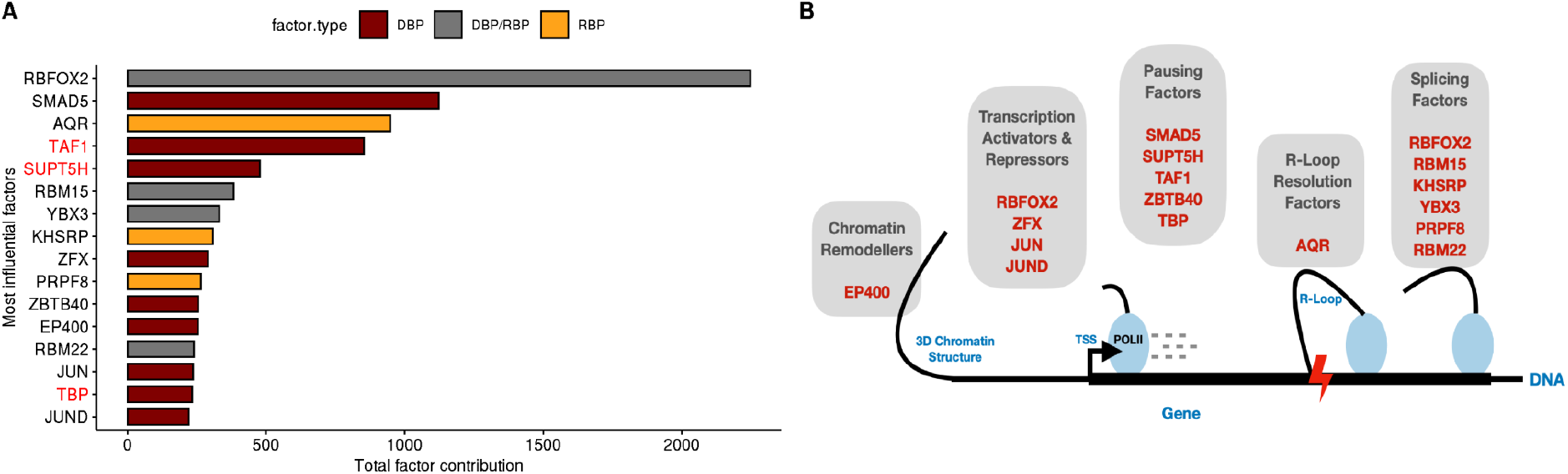
**(A)** Increasingly ordered aggregate factor contributions of factors that makeup at least 50% of model contributions. Established pausing/elongation factors are colored red. The bar fill colors identify DNA-binding (DBP; dark red), RNA-binding (RBP; orange) or DNA- and RNA-binding (DBP/RBP; grey) factors. **(B)** Functional roles of identified factors.

A minimal model that only operates on the features of these 16 most influential factors (including gene annotation and composition features) which includes only five known pausing or 7SK related factors (AQR, BRD4, SUPT5H, TAF1, TBP) achieves an R^2^ of 0.65 (on 50% holdout test data set; see **Supplementary Figure S15 & S16** performances of minimal models per cell line) and thus performs almost equally well as the full model with all 398 factors and an R^2^ of 0.68. Additionally, it outperforms the *Elongation+7SK* model (Fig. 3) which incorporates almost twice as many factors (n=27) factors of 7SK associated and established elongation factors which, although highly predictive, only achieved an R^2^ of 0.61 as compared to an R^2^ of 0.65 of the minimal model which indicates that not all pausing related factors were captured in the *Elongation+7SK* set. The minimal model (n=9) of the HepG2 data consisted of RBFOX2, AQR, TAF1, TBP, RBM15, RBM22 KHSRP, PRPF8 and YBX3, which are all included in the minimal model identified in K562.

Upon investigation of the identified most influential pausing factors (n=16, K562) defined by our model the interconnection of pausing with other RNA-processing events becomes further apparent. An interesting picture emerges considering the functional background of these factors (Fig. 4B).

### Pausing factors

Several pausing factors are well established (TAF1, TBP, SUPT5H) and occupy high ranks in our models. TAF1 and TBP are components of the pre-initiation complex (PIC). Its formation inherently leads to pausing (58). This behaviour can be modulated by other pausing factors, especially the protein complexes NELF and DSIF (SUPT5H) increase pausing whereas the P-TEFb complex associates with pause release.

### Chromatin remodelers

The chromatin remodeler EP400 had a large impact on our model. Chromatin state is defined by nucleosome positioning and posttranslational modification of its histones. It is tightly linked to transcription initiation, elongation and co-transcriptional splicing and can be actively modulated by chromatin remodelers (81–84). EP400 is a histone acetyltransferase and promotes gene activation after PIC assembly through the depositioning of H3.3/H2.AZ into promoters and enhancers (85). It interacts with the well known pausing factor MYC (26, 85, 86) and might be linked to transcriptional pausing through this association. In fact, regulation of Pol II pausing at promoter proximal nucleosomes by chromatin remodelers like for instance Chd1 (87) are already established.

### Transcriptional repressors and activators

Among the top influential factors we can find activating transcription factors ZFX, JUN and JUND as well RBFOX2 as a repressive transcription factor. ZFX family members exert a transcription activating function in multiple types of human tumors and bind downstream from the TSS at the majority of CpG island promoters regulating genes for essential housekeeping functions. ZFX family members have been suggested to act in a similar manner as the MYC family of transcription factors due to their shared pervasive binding at promoter sites as well as similar profound proliferation defects upon knockdown (88, 89). Given that MYC plays an important role in transcriptional pause release through the recruitment of P-TEFb (26, 90), a similar connection could exist for ZFX. Moreover, a comparison of the binding patterns of ZFX with Pol II and H3K4me3 have shown that ZFX is slightly downstream from the most frequent Pol II pause site and slightly upstream of the downstream peak of H3K4me3 signal (88, 89), further suggesting a role of ZFX in regulating Pol II pausing.

JUN and JUND are subcomponents of the activating protein 1 (AP-1) (91, 92) which in turn controls cell proliferation, neoplastic transformation, apoptosis and the expression of immune mediators. AP-1 is suppressed by the negative elongation factor NELF (93), but so far no regulation of transcriptional by AP-1 has been reported.

RBFOX2 acts both, as a regulator of alternative splicing as discussed later, and transcriptional repressor through the binding to chromatin-associated RNA, especially promoter-proximal nascent RNA, through the recruitment of the polycomb-complex 2 (PRC2) to its site of action (88, 94, 95). In fact, *RBFOX2* knockout cardiomyocytes were linked with decreased pausing indices and a coordinated transcriptional pause enhancing role of RBFOX2 and PRC2 at gene promoters has been suggested (95).

### Co-Transcriptional splicing and mRNA regulatory factors

The presence of several splicing associated factors (RBFOX2, PRPF8, RBM15, RBM22, KHSRP, YBX3, AQR) further strengthens the intricate connection to co-transcriptional splicing events (32, 71, 96, 97). Co-transcriptional splicing of pre-mRNAs is dependent on the availability of the nascent RNA that forms during the transcriptional cycle which in turn is a function of Pol II pausing. In fact, it has been shown that active spliceosomes are complexed to the Pol II S5P C-terminal-domain during elongation and co-transcriptional splicing (98). In particular it has also been shown that transcription kinetics strongly impact splicing decisions, such that slow Pol II elongation rates allow more time for spliceosome assembly and thereby favor splicing. Moreover, the inhibition of the spliceosomal U2 snRNP function has been shown to enhance Pol II pausing in promoter-proximal regions, to impair the recruitment P-TEFb and thereby reduce Pol II elongation velocity at the beginning of genes (79). These indicated that the release of paused Pol II requires the formation of functional spliceosomes and that a positive feedback from the splicing machinery to the transcription machinery exists. In this context, RBFOX2 acts as a well established regulator of alternative splicing (99–101) with an integral role in transcriptional pausing (95). Likewise, RBM15 (102), RBM22 (103, 104), PRPF8 (105), KHSRP (106) and YBX3 (107) as pre-mRNA splicing factors or spliceosome components are likely to have a similar connection to pausing as is the case for RBFOX2 and splicing in general.

AQR is a high ranking R-loop resolution factor (108). R-loops are RNA/DNA structures in which nascent RNA anneals back to the template DNA (109–112). It has also been suggested that R-loop formation is likely part of the mechanism for Pol II pausing (111) to hold back elongation of Pol II (113) and the DNA replisome (114). The importance of splicing events for pausing is further strengthened by splice defect induced R-loop formations as a result of increased RNA-DNA hybrid annealing due to the lack of splicing dependent nascent RNA processing which would otherwise prevent the formation of such structures through timely splicing events.

### Novel pausing factors

For the factors ZBTB40 and SMAD5 not previously associated with the regulation of pausing we suggest a novel link. ZBTB40 is not well characterized but has been established to be a regulator of osteoblast activity and bone mass (115). SMAD5, together with other SMAD proteins, is a signal transducer and is activated in the cytoplasm and accumulated in the nucleus where it regulates transcription via remodeling of the chromatin architecture through the recruitment of a variety of coactivators and corepressors to the chromatin (88, 94), suggesting a role regulating transcriptional pausing outcomes through a series of chromatin remodeling events and recruitment of transcription factors.

## DISCUSSION

The understanding of promoter-proximal Pol II pause regulatory elements is an important step towards disentangling the gene regulatory mechanisms underlying cell homeostasis and plasticity. We improved our understanding by training machine learning models that predict the extent of promoter proximal pausing from large scale genome and transcriptome binding maps, as well as gene annotation and sequence composition features providing insights into cis- and trans-acting regulatory elements underlying transcriptional pausing. Our model achieves high predictive accuracy (R^2^ ~ 0.68 with n=389, factors; R^2^ ~ 0.65 with only n=16 factors), indicating that the binding of identified trans-acting protein factors to DNA and RNA explain a large part of the variability of the extent of pausing. The accurate prediction of differential pausing based on cross cell type specific binding data (R^2^ ~ 0.52) demonstrated that the model learned general rules, which are not cell type specific. This is in line with the observation that pausing of genes is consistent across a large proportion of cell types (12). Models built from subsets of proteins implicated in all steps of gene expression, including chromatin remodelling, transcription initiation, elongation, splicing and further downstream transcript processing demonstrated high predictive power. This confirms the intimate cross talk between these processes (8, 16, 49, 66, 67, 70, 116, 117). Of note, factors implicated in splicing have the highest predictive power for pausing. This is in line with many studies that show dual roles for individual proteins such as RBFOX2 (99–101), SRSF2 (32), U2AF65 (79) or MAGOH (79) providing a direct causal link between the two processes. One important goal of our analysis was to identify novel potential pausing regulators. We achieved this using two approaches. First, we identified novel 7SK binding RBPs and showed that their binding patterns are highly predictive of pausing. Second, we analysed the feature importance in our model and pinpointed protein factors with higher feature importance than established pausing factors. Many of these factors such as RBOFX2 (99–101), AQR (108), JUN and JUND (91) have been demonstrated to affect pausing or are implicated in processes that have already been associated with pausing. These factors constitute interesting targets for further experimental validation, as our results already provide some initial mechanistic hypotheses.

We chose to analyse data from the HepG2 and K562 cell lines, since they have been extensively characterized in the ENCODE project. The number of DNA and RNA binding maps available is unparalleled and enables identification of previously unknown regulators of promoter proximal pausing. These data sets come with the limitation that not all previously characterized regulators of pausing are available. The second limitation is that only GRO-seq data and similar variations are available to quantify promoter proximal pausing. Recent multi-omics approaches based on TT-seq (118) and mNET-seq (7, 8, 119) have been applied to K562 and Raji B cell lines to estimate the kinetic rates of initiation and pause duration more precisely. These approaches provide ground for future studies of transcriptional pausing with greater precision and detail once broadly available across cell lines which would enable elaborate validation procedures. Unfortunately such data are not available for a second ENCODE cell line such that a cross validation of the model would not be possible. Taken together, our work provides a framework to further our understanding of the regulation of the critical early steps in transcriptional elongation. We expect further improvements with better kinetic profiling of the polymerase and increasing availability of binding maps or improved prediction of binding sites from sequence.

## AVAILABILITY

The code is available at https://github.com/heiniglab/POLII_pausing. All data and results are also available at 10.5281/zenodo.5236311.

## ACCESSION NUMBERS

GRO-seq data for the K562 and HepG2 cell lines were obtained from studies with GEO accessions ***GSM1480325*** and ***GSM2428726***, respectively. RNA-seq data transcript quantifications data sets (tsv-files) were taken from ENCODE from the experiment **ENCSR885DVH** with accession numbers of replicated experiments **ENCFF424CXV** and **ENCFF073NHK** for the K562 cell line, as well as the experiment **ENCSR181ZGR** with accession numbers of replicated experiments **ENCFF205WUQ, ENCFF915JUZ** for the HepG2 cell line. ENCODE Accession number of CHIP-seq and eCLIP-seq data sets can be found in supplementary tables S3 & S4 and S7 & S8, respectively. Annotations of housekeeping genes were taken from (54) (see **Supplementary Table S21;** housekeeping.RDS in zenodo repository). CpG island annotations were taken from the UCSC golden path for the hg19 genome build (cpgIslandExt.txt.gz) (see **Supplementary Table S21;** cpg.islands.RDS in zenodo repository). Gene annotations along with HGNC and RefSeq metadata files were taken from GENCODE (see **Supplementary Table S21**). CAGE transcription start sites are provided in the zenodo repository as an R-data structure (CTSS.RDS).

## ACKNOWLEDGEMENT

We would like to acknowledge the ENCODE Consortium and the ENCODE production laboratories generating the particular datasets. We would like to acknowledge insightful discussions with Dr. Jernej Murn.

## SUPPLEMENTARY DATA

Supplementary Data are available at NAR online:

Supplementary material (figures and tables as pdf)

Supplementary tables (as xlsx).

