## Supplementary Material for "Predictive model of transcriptional elongation control identifies trans regulatory factors from chromatin signatures"

* Corresponding author

**SUPPLEMENTARY FIGURES**

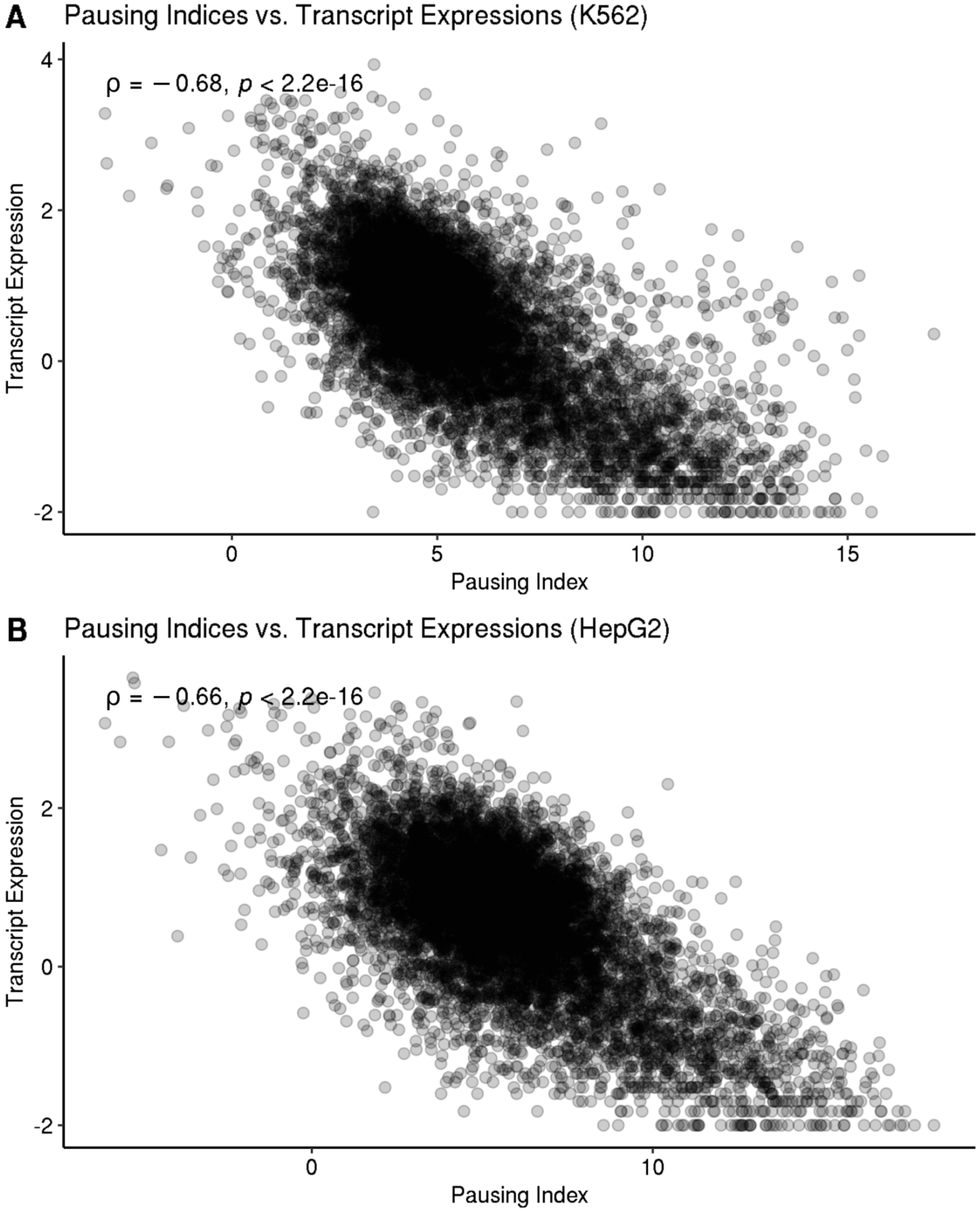


**Supplementary Figure 1: Pausing Index Optimization.** Inverse correlation of pausing indices (x-axis) with transcript expressions (FPKMs, y-axis) in the K562 (A) and HepG2 (B) cell line. Pearson's correlation coefficient rho (ρ) with the associated p-value is depicted in the upper left.

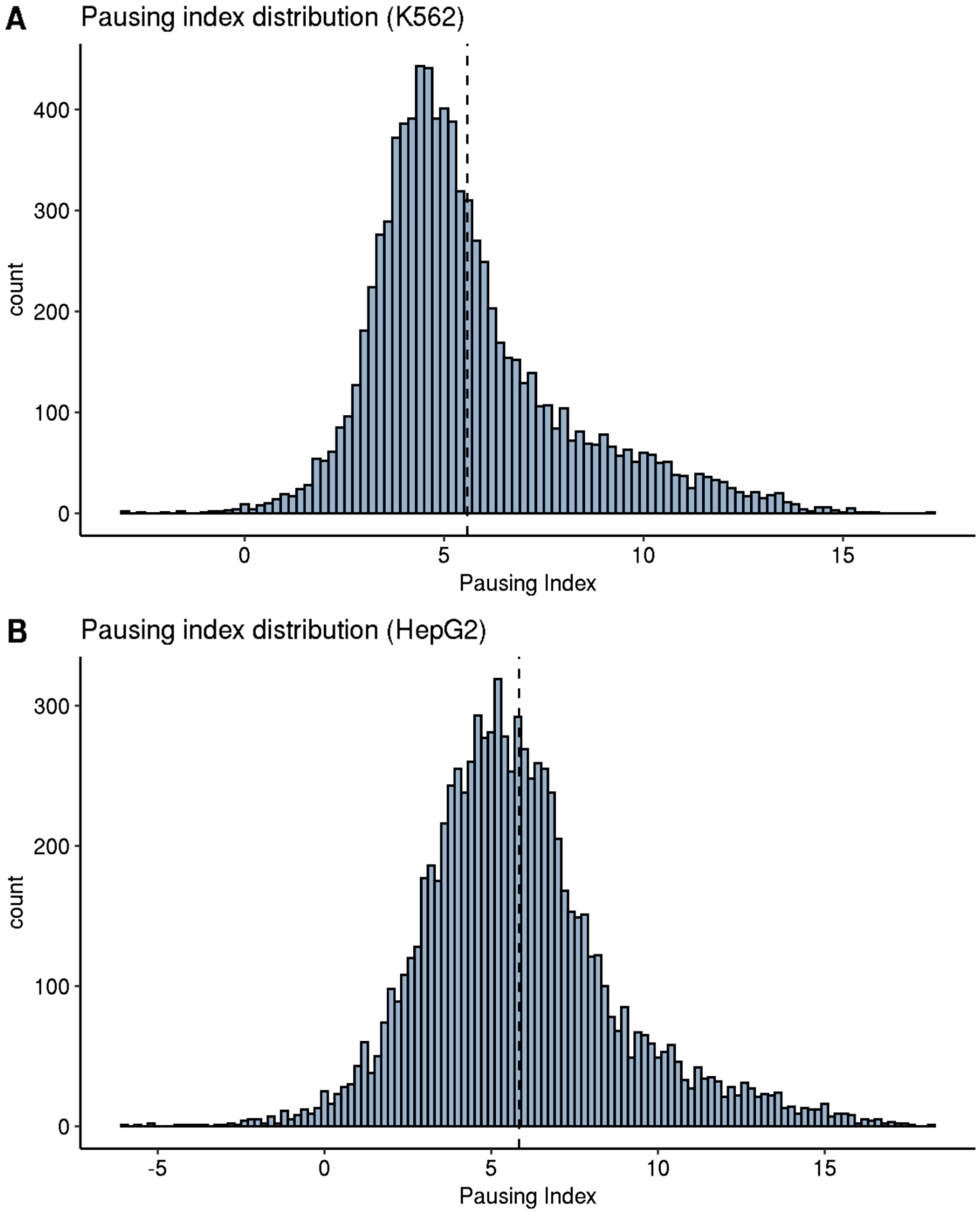


**Supplementary Figure 2: Pausing Index Distribution.** Histograms of the distribution of pausing indices (PIs) in the K562 (A) and HepG2 (B) cell line. Dashed lines indicate the mean pausing indices, the x-axes the PIs and the y-axes the PI counts.

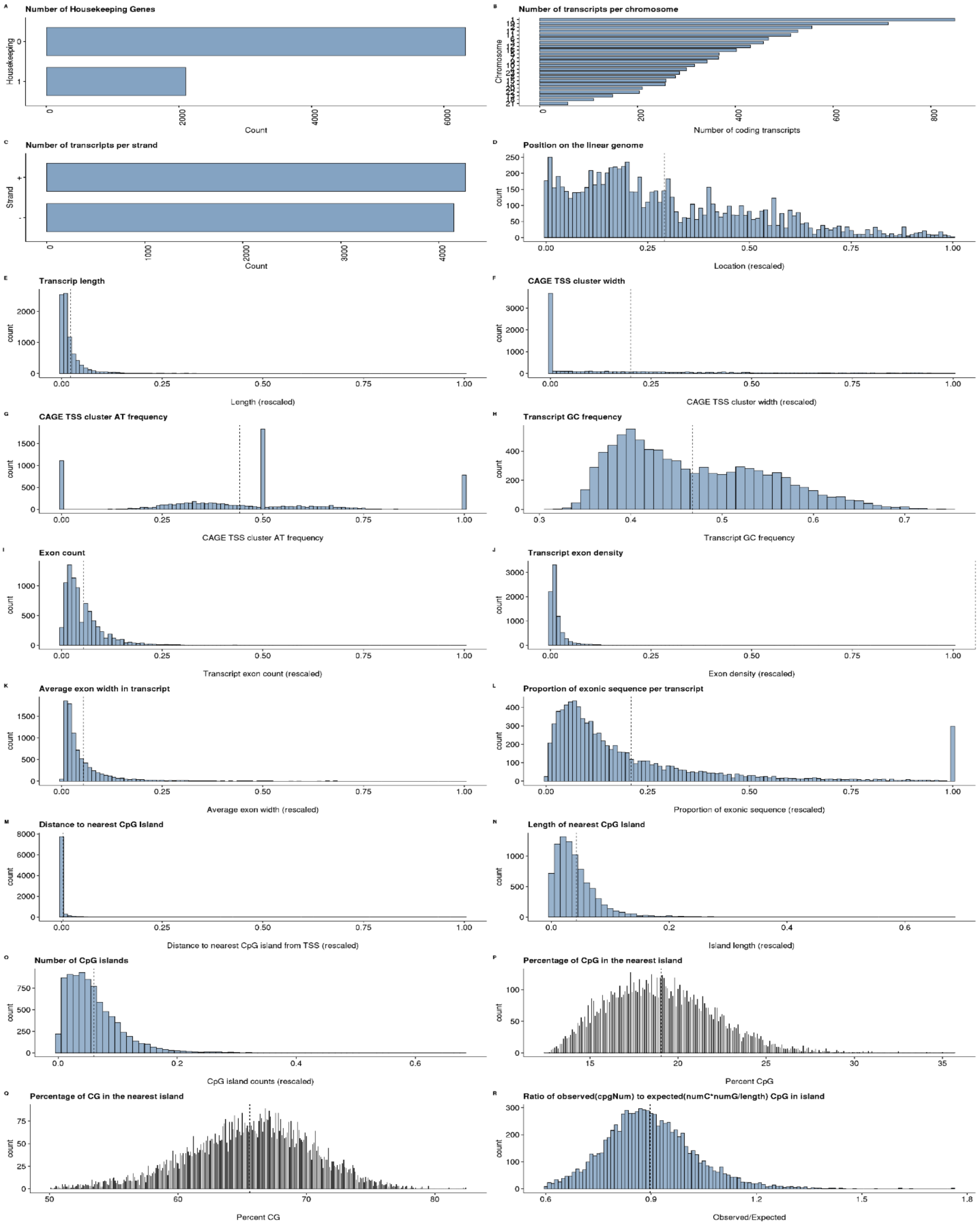


**Supplementary Figure 3: Gene annotation and composition features (K562).** Distribution of gene annotation and sequence composition features in the K562 cell line. Numeric features were rescaled to the range [0;1]. In sub-figures A-C the x-axes show the counts of features and the y-axes the feature values. In sub-figues D-R the x-axes show the feature values and the y-axes the counts of features.

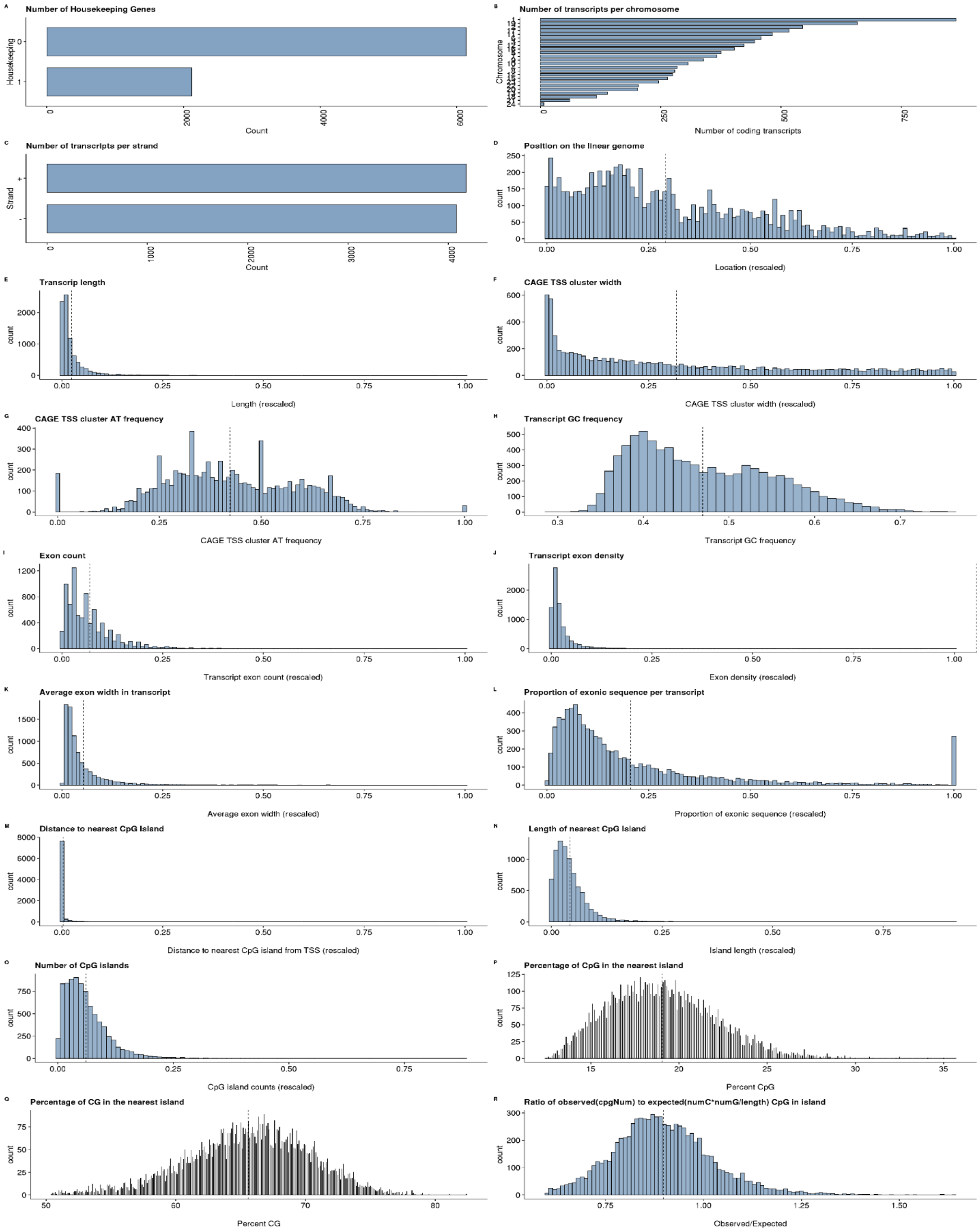


**Supplementary Figure 4: Gene annotation and composition features (HepG2).** See caption of supplementary figure 3 for more details.

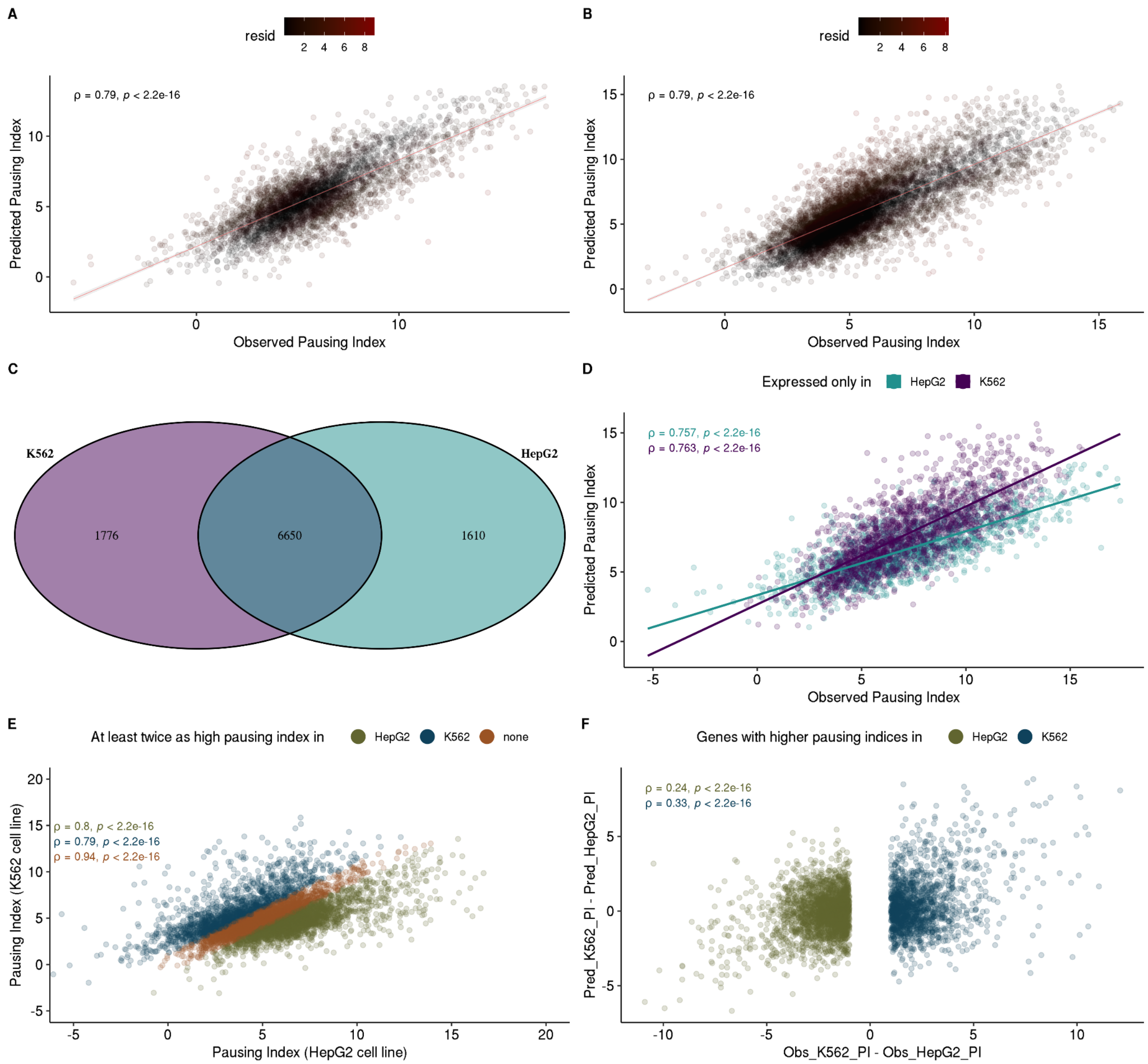


**Supplementary Figure 5: Figure 2 analog for the HepG2 cell line.** See caption of main figure 2 for more details.

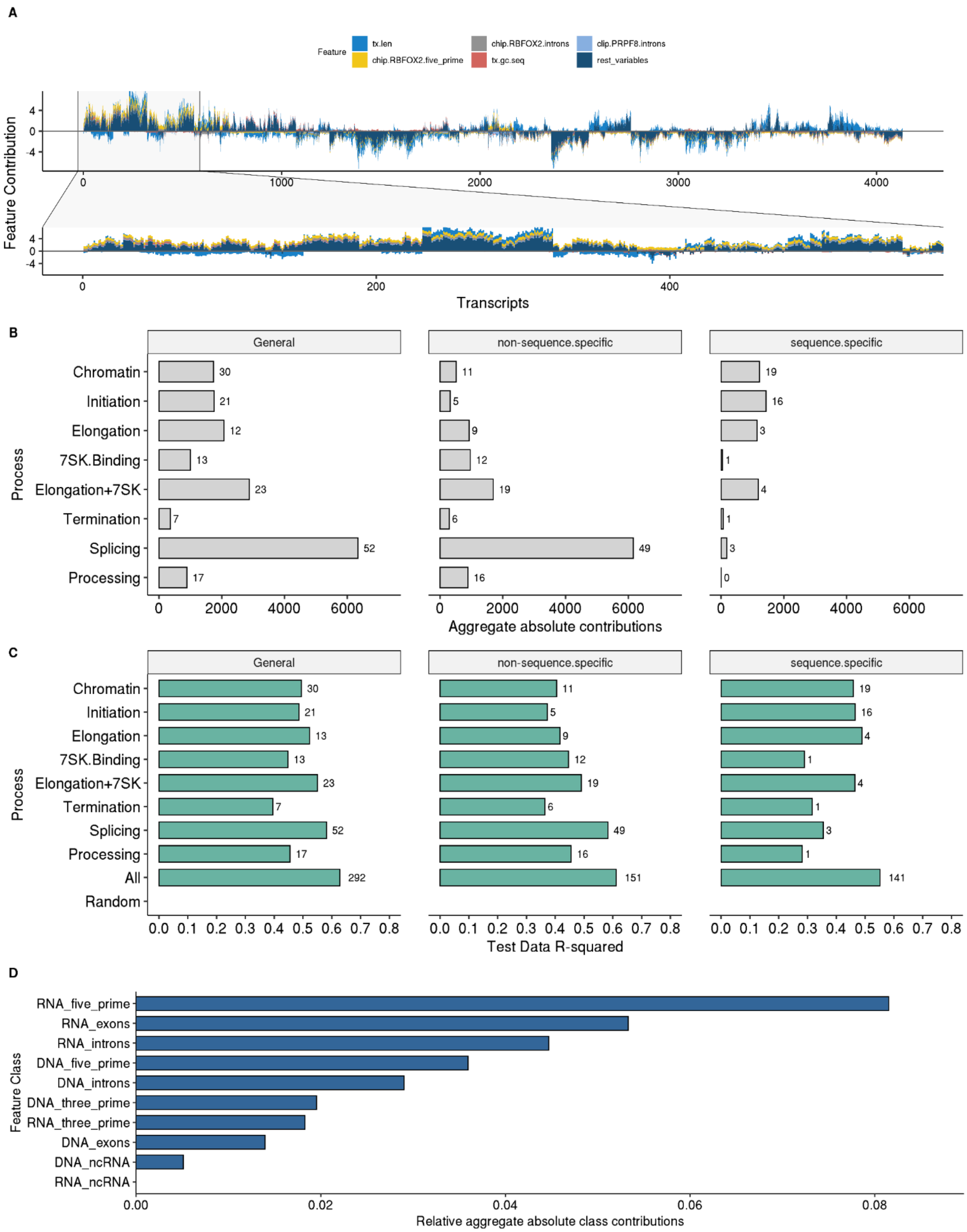


**Supplementary Figure 6: Figure 3 analog for the HepG2 cell line.** See caption of main figure 3 for more details.

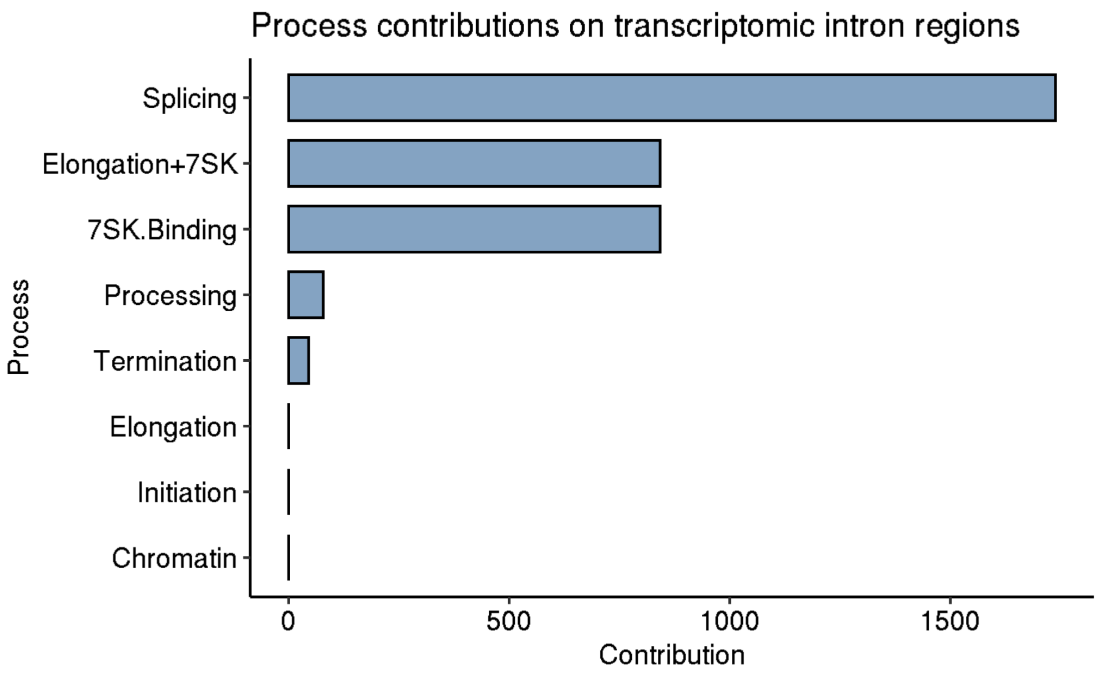


**Supplementary Figure 7: Feature contributions on RNA introns (K562 cell line).** Aggregate feature contributions (x-axis) of RNA intron binding factors by functional classes (y-axis).

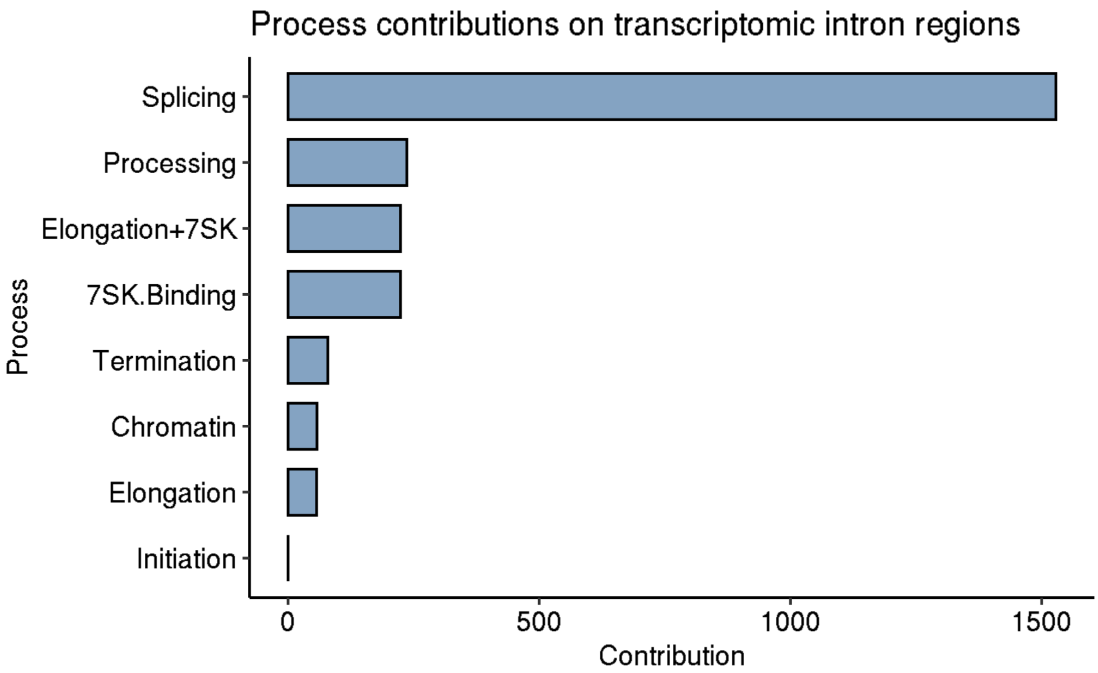


**Supplementary Figure 8: Feature contributions on RNA introns (HepG2 cell line).** See caption of supplementary figure 7 for more details.

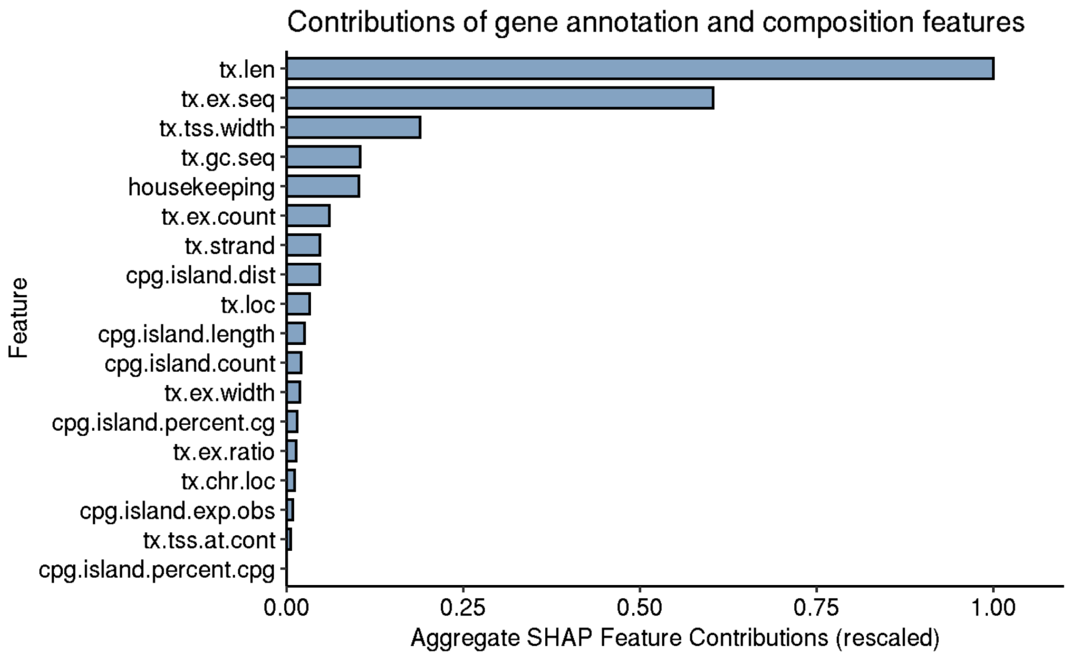


**Supplementary Figure 9: Aggregate feature contributions of gene annotation and composition features (K562).** Aggregate feature contributions (x-axis) of gene annotation and composition features (y-axis) in the K562 cell line.

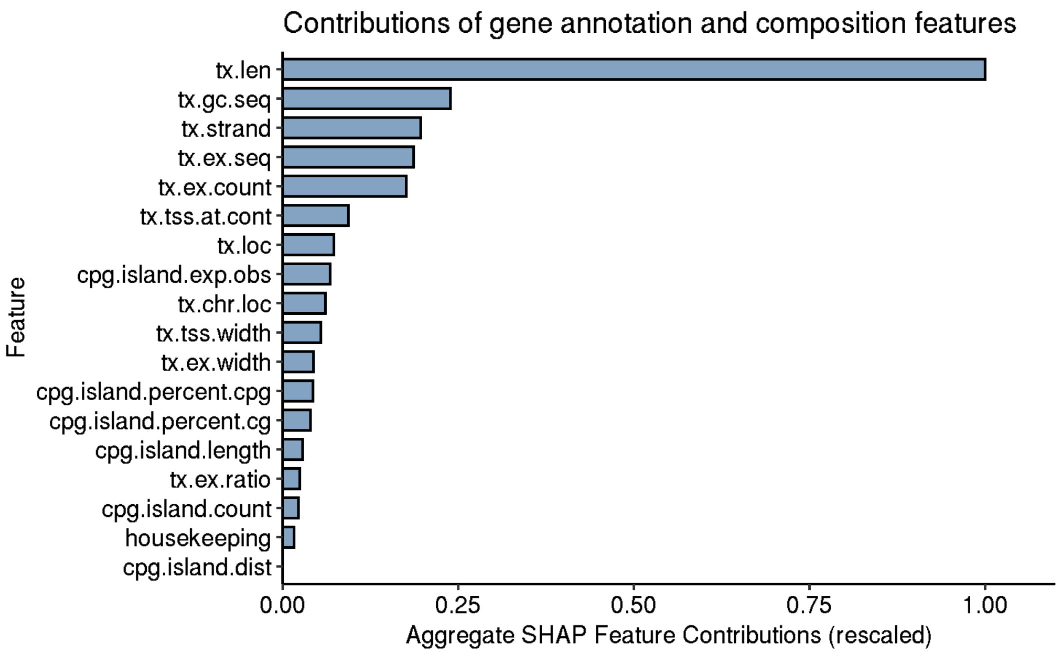


**Supplementary Figure 10: Aggregate feature contributions of gene annotation and composition features (HepG2).** See caption of supplementary figure 9 for more details.

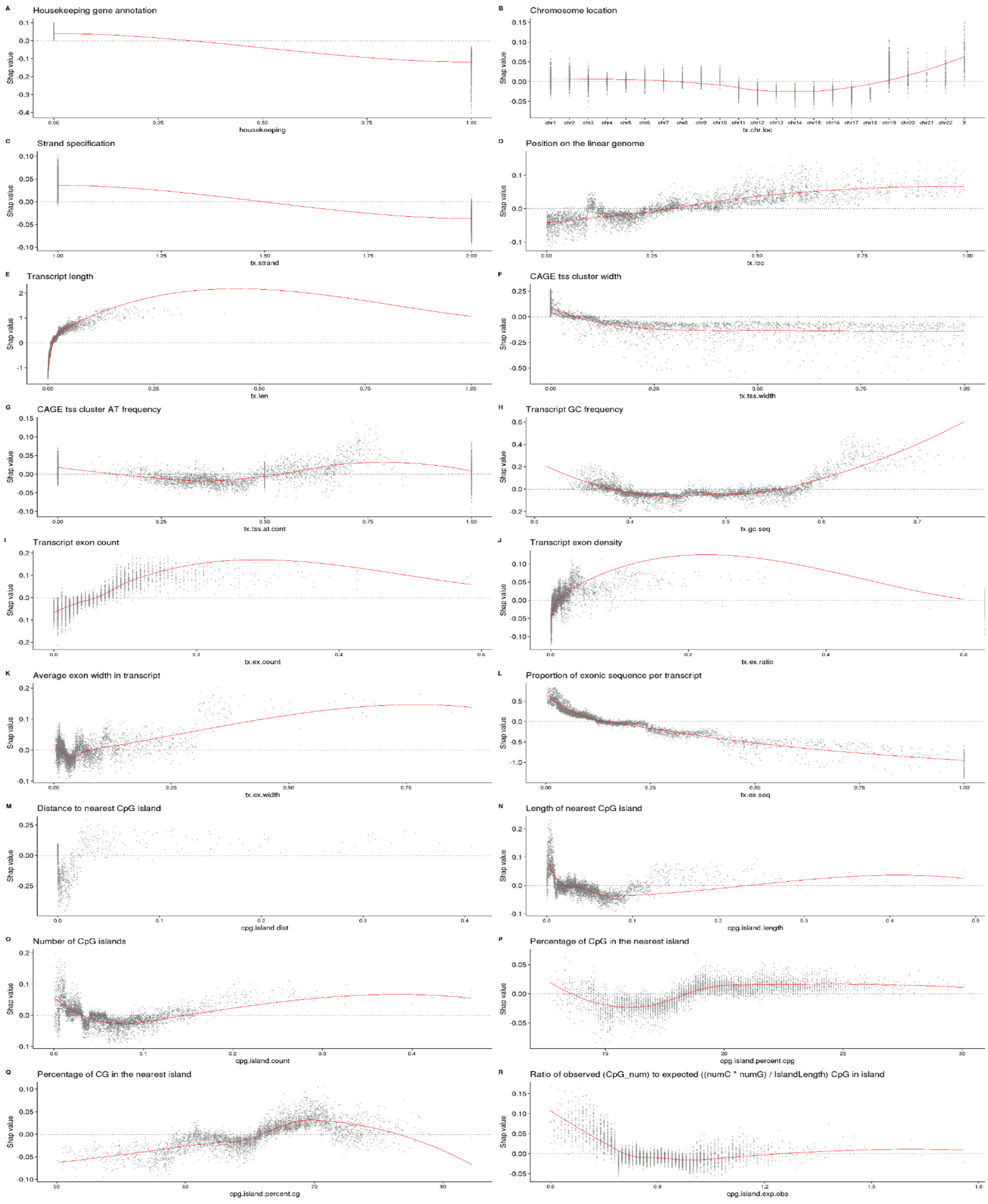


**Supplementary Figure 11: Model feature contribution distributions of gene annotation and composition features (K562).** Feature contributions (y-axes) of gene annotation and composition features values (x-axes) in the K562 cell line. For “tx.strand” feature, “1” denotes “+” (forward) strand and “2” denotes “-” (reverse strand).

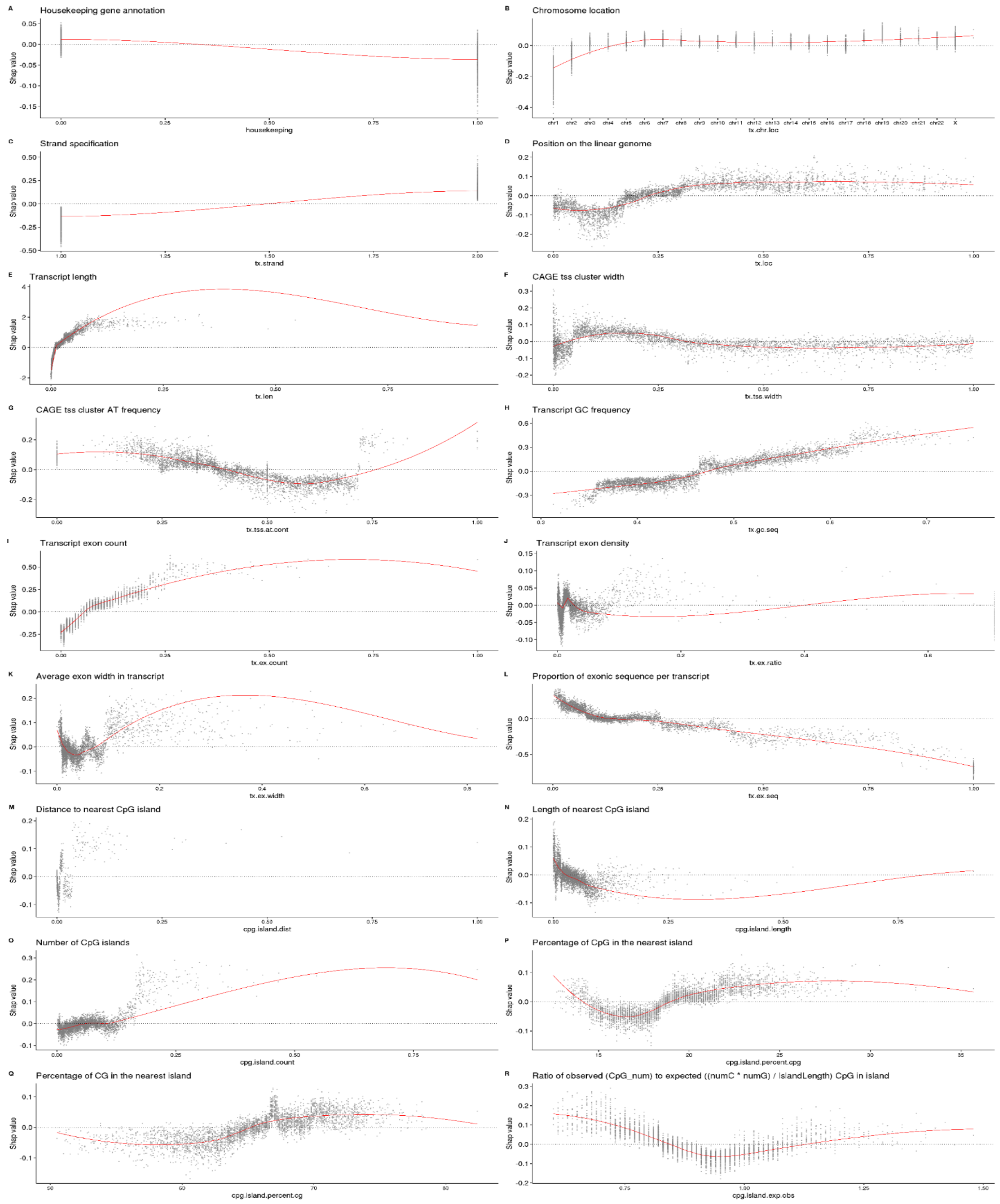


**Supplementary Figure 12: Model feature contribution distributions of gene annotation and composition features (HepG2).** See caption of supplementary figure 11 for more details.

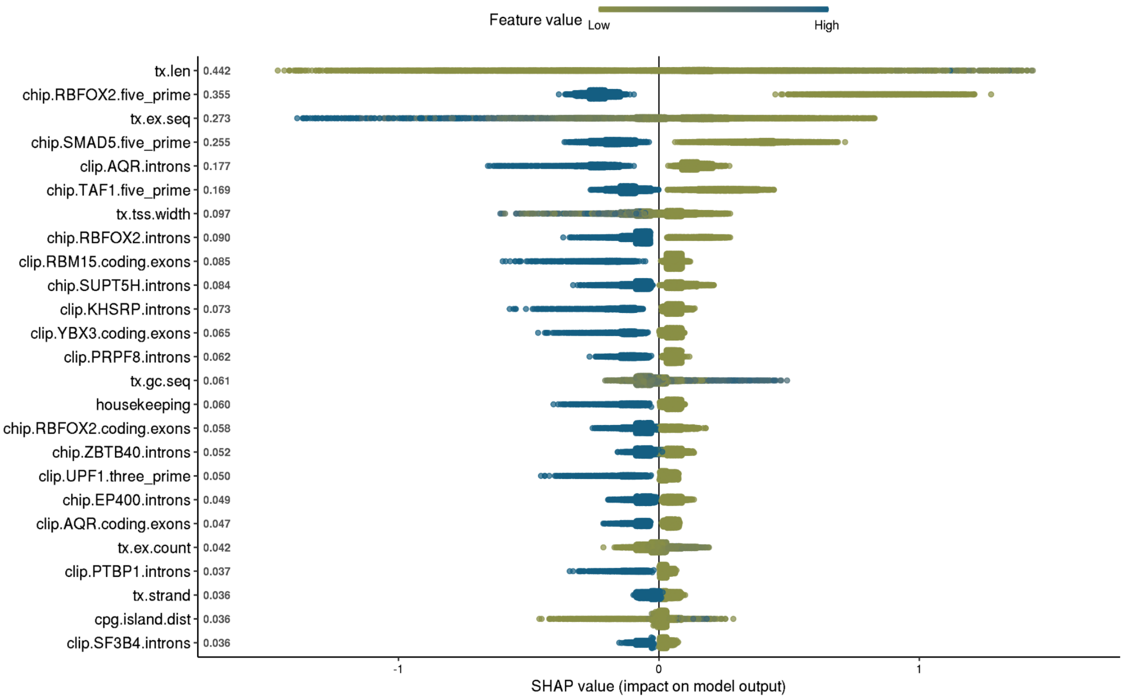


**Supplementary Figure 13: Feature Contributions (K562).** Feature contribution (x-axis) of the top 25 features (y-axis) from the full (*All)* individual K562 model.

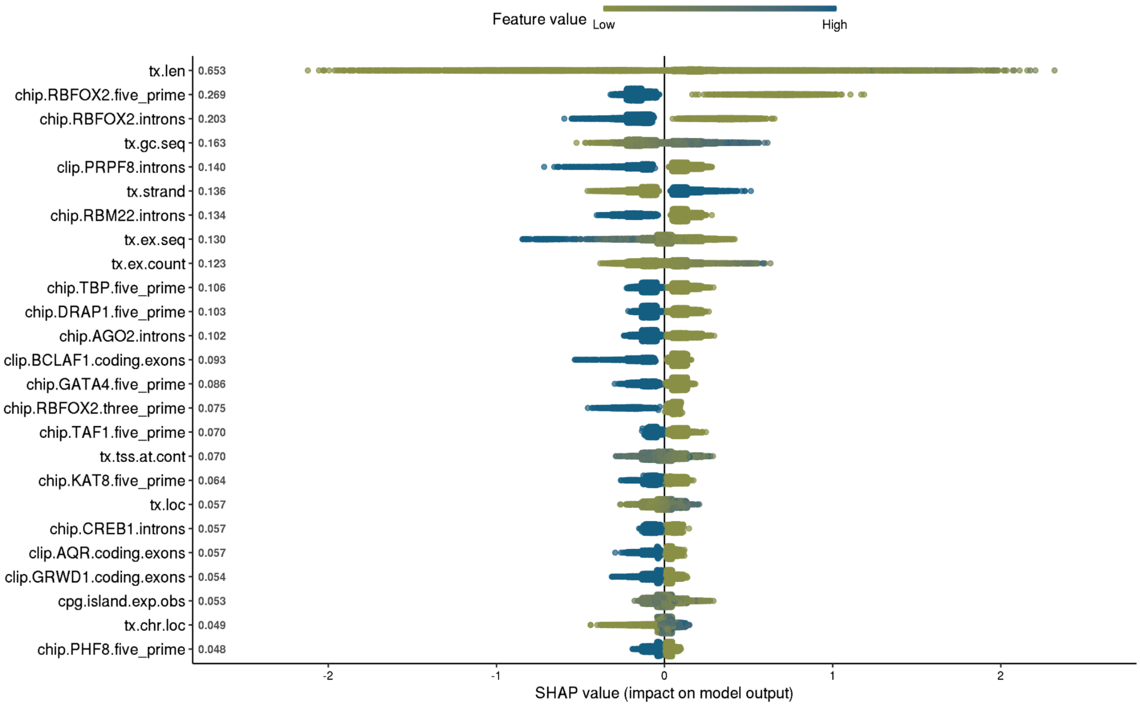


**Supplementary Figure 14: Feature Contributions (HepG2).** See caption of supplementary figure 13 for more details.

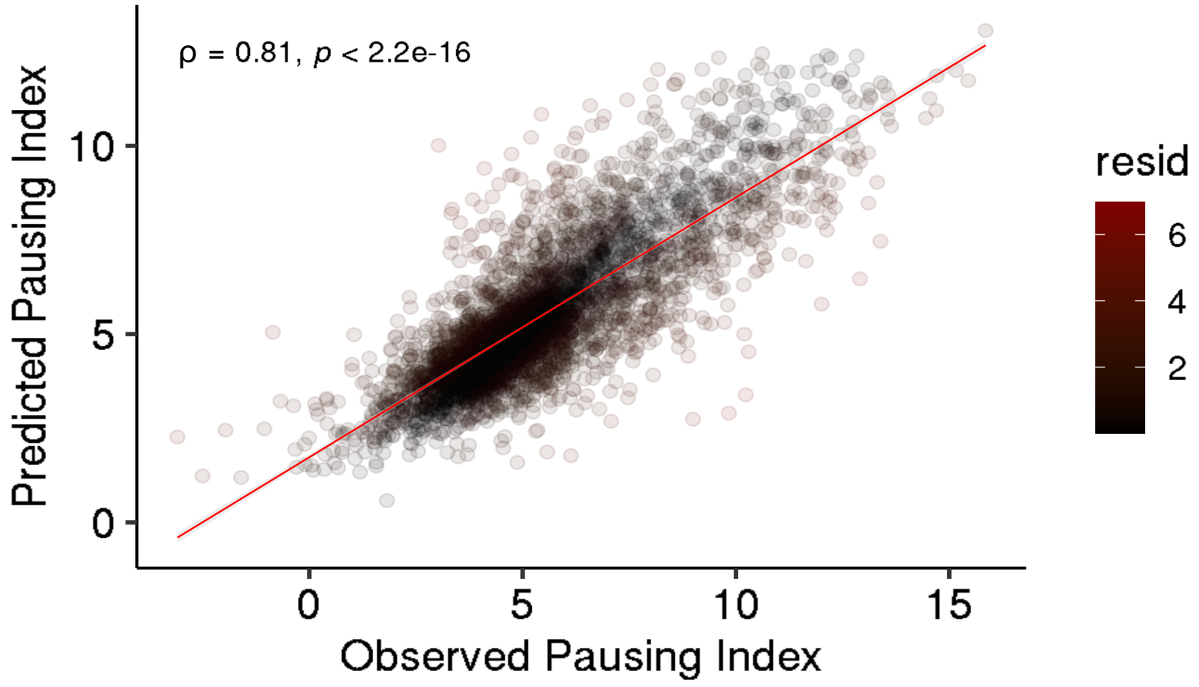


**Supplementary Figure 15: Minimal model performance.** Observed (x-axis) vs predicted (y-axis) pausing index of the 16 most influential factor model for the K562 cell line.

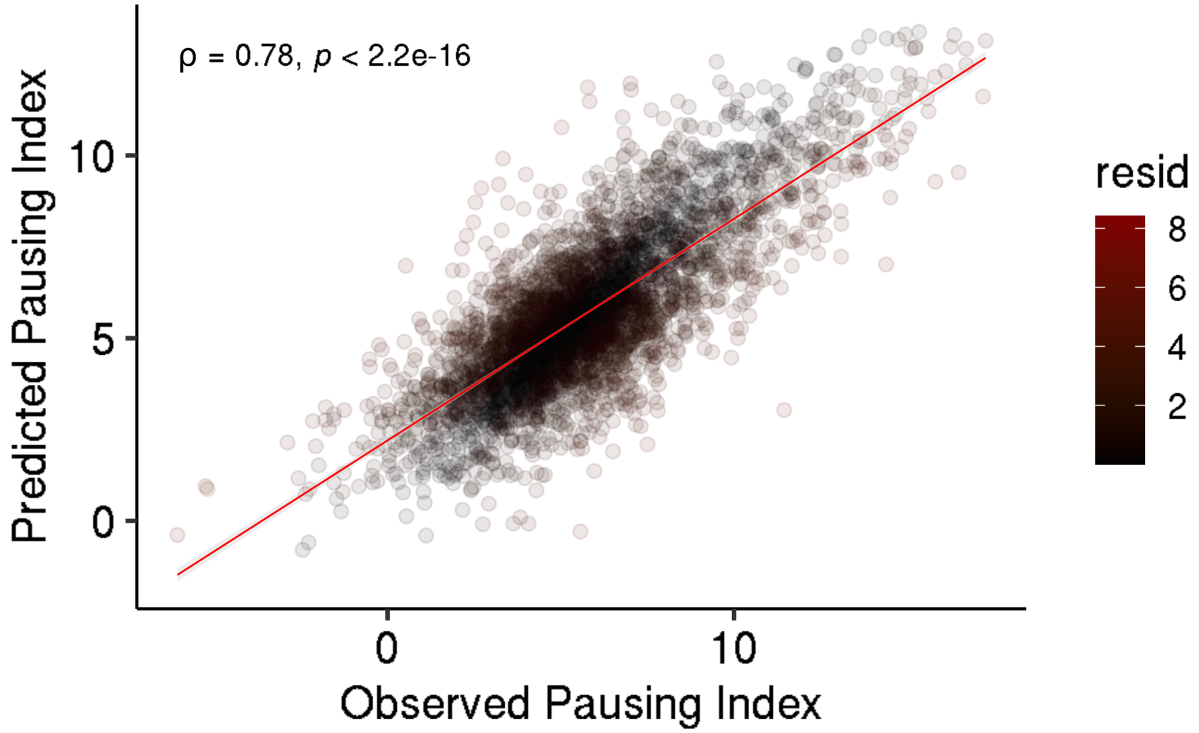


**Supplementary Figure 16: Minimal model performance.** Observed (x-axis) vs predicted (y-axis) pausing index of the 9 most influential factor model for the HepG2 cell line.

### SUPPLEMENTARY TABLES

**Supplementary Table S1 (see xls file sheet "S1 K562 CHIPseq Factors") :** List of factors from the ENCODE CHIP-seq experiments for the K562 cell line.

**Supplementary Table S2 (see xls file sheet "S2 HepG2 CHIPseq Factors") :** List of factors from the ENCODE CHIP-seq experiments for the HepG2 cell line.

**Supplementary Table S3 (see xls file sheet "S3 K562 CHIPseq Accessions") :** List of ENCODE CHIP-seq experiment accession numbers for the K562 cell line.

**Supplementary Table S4 (see xls file sheet "S4 HepG2 CHIPseq Accessions"):** List of ENCODE CHIP-seq experiment accession numbers for the HepG2 cell line.

**Supplementary Table S5 (see xls file sheet "S5 K562 eCLIPseq Factors"):** List of factors from the ENCODE eCLIP-seq experiments for the K562 cell line.

**Supplementary Table S6 (see xls file sheet "S6 HepG2 eCLIPseq Factors"):** List of factors from the ENCODE eCLIP-seq experiments for the HepG2 cell line.

**Supplementary Table S7 (see xls file sheet "S7 K562 eCLIPseq Accessions") :** List of ENCODE eCLIP-seq experiment accession numbers for the K562 cell line.

**Supplementary Table S8 (see xls file sheet "S8 HepG2 eCLIPseq Accessions"):** List of ENCODE eCLIP-seq experiment accession numbers for the HepG2 cell line.

**Supplementary Table S9 (see xls file sheet "S9 K562 7SK Binding Factors"):** List of factors that bind the 7SK ncRNA in the K562 cell line. Please note that binding signals of pseudo 7SK ncRNA transcript variants expressed above median ncRNA expression levels were included. Their consideration was supported by the transcripts' high mean pairwise sequence similarity (41) of 0.74 and high mean conservation score of 923.58 (PAM250 scoring matrix) resulting from a multiple sequence alignment (ClustalW alignment) of corresponding 7SK transcripts.

**Supplementary Table S10 (see xls file sheet "S10 HepG2 7SK Binding Factors"):** List of factors that bind the 7SK ncRNA in the HepG2 cell line. Please note that binding signals of pseudo 7SK ncRNA transcript variants expressed above median ncRNA expression levels were included. Their consideration was supported by the transcripts' high mean pairwise sequence similarity (41) of 0.81 and high mean conservation score of 302.29 (PAM250 scoring matrix) resulting from a multiple sequence alignment (ClustalW alignment) of corresponding 7SK transcripts.

**Supplementary Table S11 (see xls file sheet "S11 K562 Factor Bindings"):** Number of bindings on genomic and transcriptomic transcript regions per factor in the K562 cell line.

**Supplementary Table S12 (see xls file sheet "S12 HepG2 Factor Bindings"):** Number of bindings on genomic and transcriptomic transcript regions per factor in the HepG2 cell line.

**Supplementary Table S13 (see xls file sheet "S13 Known Pausing Factors"):** List of known pausing factors from the literature.

**Supplementary Table S14 (see xls file sheet "S14 K562 Factors per Process"):** List of factors in the K562 cell line per functional process.

**Supplementary Table S15 (see xls file sheet "S15 HepG2 Factors per Process"):** List of factors in the HepG2 cell line per functional process.

**Supplementary Table S16 (see xls file sheet “S16 K562 Sequence Specificity”):** An indicator matrix whether a factor in the K562 cell line is sequence specific (column *SS*), non-sequence specific (column *NSS*), a RNA-binding factor (column *RBP*) or a DNA-binding factor (column DBP).

**Supplementary Table S17 (see xls file sheet “S17 HepG2 Sequence Specificity”):** An indicator matrix whether a factor in the HepG2 cell line is sequence specific (column *SS*), non-sequence specific (column *NSS*), a RNA-binding factor (column *RBP*) or a DNA-binding factor (column DBP).

**Supplementary Table S18 (see xls file sheet "S18 Subspace Factors Presence"):** An indicator matrix whether a factor was present in any of the feature subspaces. “1” denotes present, “0” denotes” not present

**Supplementary Table S19 (see xls file sheet “S19 Hyperparameters”):** Specification of hyperparameters of the Extreme Gradient Boosting Tree regressor.

**Supplementary Table S20 (see xls file sheet “S20 All Model Results”):** Model results for each cell line and each feature subspace. Column *subspace* gives the feature subspace the model was trained on. An appendix of “*ss*” to the feature subspace name indicates a model trained on binding features of sequence specific factors and “*nss”* of non-sequence specific factors. The model type *synchronised.model.matrices* as opposed to *individual.model.matrices* refers to a model that was trained on features observed in both of the cell lines (K562 and HepG2). Column *train.rsqrd* gives the $R^2$ performance of the 5-fold cross-validation procedure. Column *test.rsqrd* gives the performance on a 50% hold out test data set taken before training. Column *mean.shap* gives the average feature contribution over all factor associated binding features.

**Supplementary Table S21 (see xls file sheet “S21 Data Accessions”):** List of data accession numbers.
